# Identification of biochemically neutral positions in liver pyruvate kinase

**DOI:** 10.1101/632562

**Authors:** Tyler A. Martin, Tiffany Wu, Qingling Tang, Larissa L. Dougherty, Daniel J. Parente, Liskin Swint-Kruse, Aron W. Fenton

**Affiliations:** Department of Biochemistry and Molecular Biology, The University of Kansas Medical Center, Kansas City, KS 66160; Department of Family and Community Medicine, The University of Kansas Medical Center, Kansas City, KS 66160

**Keywords:** Neutral position, allosteric regulation, pyruvate kinase, neutral substitutions

## Abstract

Understanding how each residue position contributes to protein function has been a long-standing goal in protein science. Substitution studies have historically focused on conserved protein positions. However, substitutions of nonconserved positions can also modify function. Indeed, we recently identified nonconserved positions that have large substitution effects in human liver pyruvate kinase (hLPYK), including altered allosteric coupling. To facilitate a comparison of which characteristics determine when a nonconserved position does vs. does not contribute to function, the goal of the current work was to identify neutral positions in hLPYK. However, existing hLPYK data showed that three features commonly associated with neutral positions – high sequence entropy, high surface exposure, and alanine scanning – lacked the sensitivity needed to guide experimental studies. We used multiple evolutionary patterns identified in a sequence alignment of the PYK family to identify which positions were least patterned, reasoning that these were most likely to be neutral. Nine positions were tested with a total of 117 amino acid substitutions. Although exploring all potential functions is not feasible for any protein, five parameters associated with substrate/effector affinities and allosteric coupling were measured for hLPYK variants. For each position, the aggregate functional outcomes of all variants were used to quantify a “neutrality” score. Three positions showed perfect neutral scores for all five parameters. Furthermore, the nine positions showed larger neutral scores than 17 positions located near allosteric binding sites. Thus, our strategy successfully enriched the dataset for positions with neutral and modest substitutions.

## Introduction

In protein structure/function studies, a common experimental design is to substitute an amino acid position and evaluate whether protein function is perturbed. Such studies often target positions that are conserved within a protein family because conserved positions are likely to have key roles in function. The contributions of nonconserved positions have historically been overlooked^1^, although their substitution can also modify function ^2,3^ and cause disease (*e.g*., ^4^). We have been using human liver pyruvate kinase (hLPYK) to probe the functional roles of nonconserved protein positions. In our studies, substitutions at nonconserved positions in and near the allosteric binding sites modulated multiple functional parameters^2,3,5-9^. To better understand how the nonconserved positions contribute to function, we wish to compare their biochemical, biophysical and structural characteristics to nonconserved positions that have little effect on function. Thus, the goal of the current study was to identify hLPYK positions that are biochemically neutral (*i.e*., those that can be substituted with a range of replacement amino acids with no change in function).

We first assessed three of the “usual” strategies to identify neutral positions, but existing data showed that these metrics lacked the sensitivity needed to guide experimental studies. Thus, we developed an alternative strategy to select positions likely to be neutral. We assessed nine of those positions using biochemical assays that report five different parameters associated with different hLPYK functions. Our approach successfully identified multiple positions in hLPYK that showed little to no functional consequences when substituted. These positions provide a useful set of biochemically neutral positions that will serve as controls in our ongoing studies.

## Materials and Methods

### Multiple sequence alignment for the PYK family and overview of analysis strategy

For the PYK family, we used the previously constructed sequence alignment with 241 representative PYK sequences from all kingdoms of life^4^. The sequences for this alignment were sampled so that no group was over-represented; in particular, only one representative was included for each of the four mammalian isozymes. Note that the mammalian isozymes differ in their allosteric regulation^10,11^, which by definition, must arise from amino acid changes at nonconserved positions. Sugar phosphates are common regulators in most known PYK isozymes, although effector specificity differs among isozymes. In addition, inhibition by amino acids is isolated to higher animal species. For this work, we assessed the PYK sequence alignment with four types of sequence analyses to locate candidate neutral positions: conservation (Shannon entropy), specificity (two-entropies analysis), co-evolution (several algorithms, below) and SNAP (neural network-based). Each of these analyses quantifies an evolutionary constraint pattern (Supplemental Figure 1). Candidate neutral positions are those that have the least match to any of these patterns.

### Conservation

To determine conservation/nonconservation for positions in hLPYK, we used the PYK sequence alignment of Pendergrass *et al*.^4^ to calculate Shannon sequence entropy with the program BioEdit^12^ using the equation

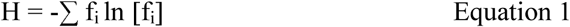

where H is the sequence entropy for a given position and f_i_ is the frequency for each of the i’th type of amino acid (Ser, Asp, Leu, *etc*.) with f_i_ ≠ 0 at that position. As with most protein families, sequence entropy calculations generate a continuum of scores for the PYK sequence alignment (*e.g*., Figure 1); no natural threshold exists to clearly demarcate “conserved” from “nonconserved” positions. We previously discussed various approaches to thresholding continuous scores^13^. However, for the current work, we do not need to identify a conservation/nonconservation threshold due to our use of the most extreme “least patterned” positions, a classification that is defined below.

**Figure 1.**
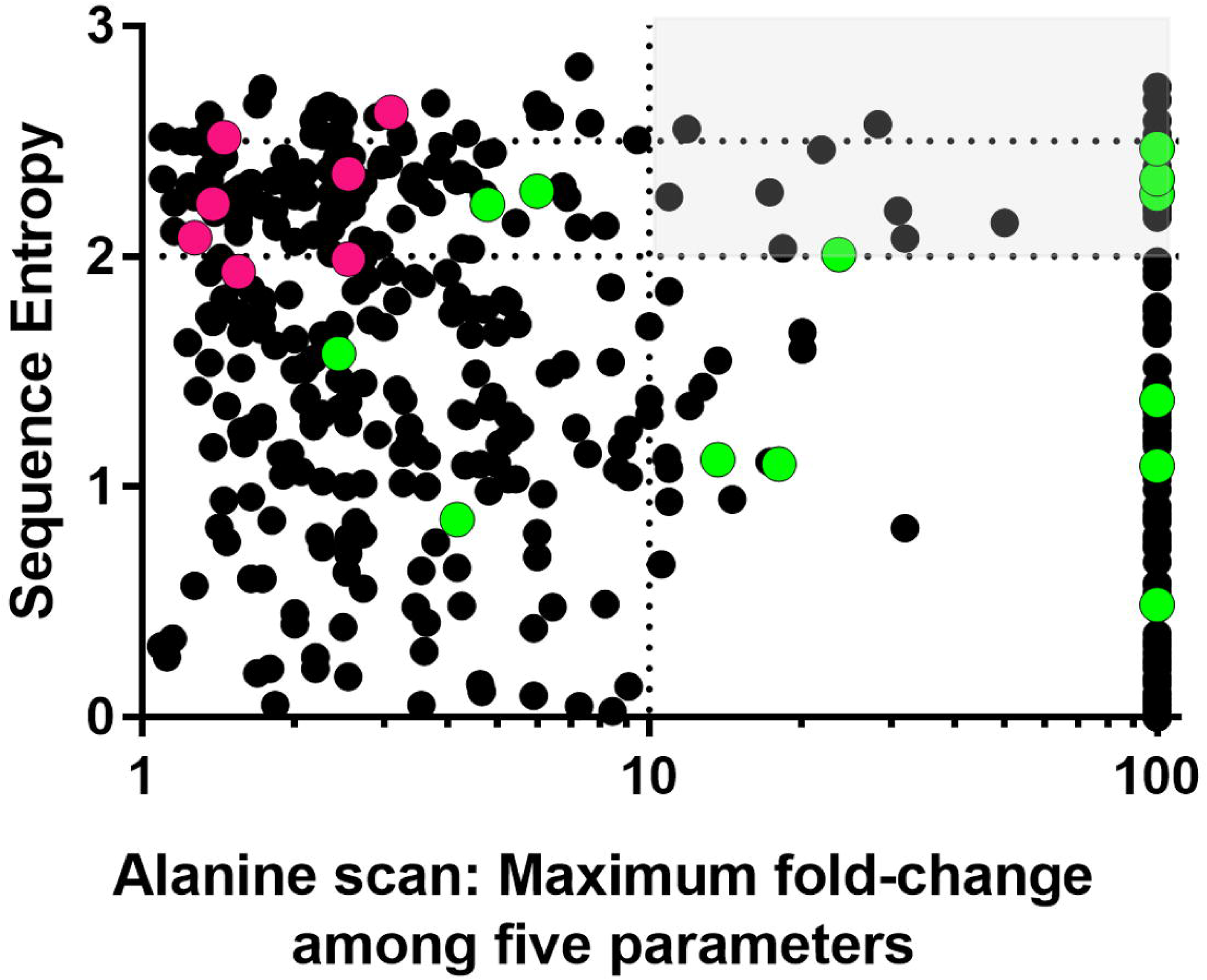
High sequence entropy does not identify functionally neutral positions in hLPYK. Using a whole-protein alanine scan of hLPYK ^9^, the maximum fold-change (increase or decrease) relative to wildtype was calculated from the five functional parameters measured experimentally. If the alanine variant abolished binding to an allosteric ligand or allosteric response, it is assigned a value of 100. If a position was alanine or glycine in wild-type PYK, or if an alanine substitution abolished all enzymatic activity, no data are shown (*i.e*., some residues are not represented on this graph). Sequence entropies were calculated from the PYK sequence alignment of Pendergrass *et al*., ^4^ (black dots). Higher sequence entropies represent lower conservation, and the dotted lines and light gray box are used to highlight non-conserved positions for which an alanine substitution leads to more than a 10-fold change in at least one functional parameter. The overlaid green dots show previously characterized positions in the allosteric sites ^2,3,5,27^ and the magenta dots show 7 of the positions substituted in this study (the other two positions included in the current study were not reported in the alanine scan).

The calculated sequence entropy scores were compared to existing hLPYK substitution datasets to show that sequence entropy alone could not identify functionally neutral positions (Figure 1). Thus, we used a range of strategies to identify patterns within the sequence alignments (detailed below) and combined these strategies to identify a set of “least patterned” nonconserved positions for experimental characterization.

### Phylogenetic and co-evolutionary patterns of change

Patterns of change in a sequence alignment can be detected by analyzing the amino acids present in each column of a sequence alignment. Several algorithms have been developed to track which amino acid changes mirror the branching of a protein families’ phylogenetic tree^14-16^. For this work, we chose to use two entropies analysis – objective (TEA-O) since it generates “conservation” and “specificity” scores for each position by measuring the conservation at different branch levels of the phylogenetic tree. Another pattern of change in sequence alignments is the pairwise “co-evolution” observed when pairs of positions are observed to change together. Dozens of co-evolutionary algorithms are now available; for many protein families, the score correlation among different algorithms is low^17,18^. Previous analyses with other protein families found that different algorithms each detected “important” protein positions^18,19^, and thus may detect different evolutionary constraints. Thus, the “least patterned” positions for this work should have low co-evolution scores in multiple co-evolutionary algorithms; we used four: McLachlan-based Substitution Correlations (McBASC)^20-22^, Observed Minus Expected Squared (OMES)^23,24^, Explicit Likelihood of Subset Covariation (ELSC)^25^, and Z-Normalized Mutual Information (ZNMI)^17^.

The PYK sequence alignment was used to calculate the TEA-O^14^ conservation and specificity scores and pairwise co-evolutionary scores from the four different algorithms (listed above) using the ensemble-average implementation in the software suite “Co-evolutionary utilities”^18^. Note that the four co-evolutionary algorithms have different mathematical bases^18^. Since the phylogenetic and four co-evolutionary algorithms utilized different scoring scales, we next used their maximum and minimum scores to re-scale all scores from 0.01 (least patterned) to 1 (most patterned). The six standardized scores were then multiplied to generate a single composite score, hereafter referred to as the “least patterned” score. Throughout the text, PYK positions with the most-significant, least-patterned score (with the exception of those excluded by SNAP below) are referred to as “least patterned positions.”

### SNAP (screening for non-acceptable polymorphisms) analysis

Using the rank-ordered list of the composite “least patterned” score, PYK positions with the most extreme scores were further assessed using SNAP^26^. This machine-learning algorithm uses a variety of predicted structural features and a few sequence features (but not co-evolution or phylogeny) to predict neutral/non-neutral outcomes for amino acid substitutions. (Note that SNAP predicts the consequences of individual amino acid substitutions, *e.g*., E277K, rather than for columns of a sequence alignment.) For the current work, all 19 possible substitutions were predicted for the PYK positions with the most extreme least patterned scores. Twenty of the top 22 positions had at most one non-neutral prediction; furthermore, the non-neutral substitutions were either a proline or tryptophan (positions 75, 138, 210, 388, and 470). Since proline has substantial effects on the backbone and tryptophan inserts a large side chain, we considered the aggregate of these SNAP outputs to be consistent with predictions of neutral positions. Two positions (59 and 248) were predicted to have multiple non-neutral substitutions and were excluded from further consideration.

### Biochemical characterization

Experimental assays, the creation of mutations, protein expression, protein preparation, and data analyses were performed the same as previously reported ^5,9,27^. In brief, variant hLPYK proteins were created by mutating the coding region of the pLC11 plasmid (originally a kind gift from Dr. Andrea Mattevi ^28^) using QuikChange (Stratagene) and primers designed to individually obtain all 19 substitutions at each position. After QuikChange mutagenesis, we sequenced a single colony from each mutagenic reaction. Although we did not obtain all 19 substitutions, this method generated a range of substitutions at each position in a cost-effective manner without introducing a human bias into the final set of substitutions.

Wildtype and mutated genes were expressed in FF50 *E. coli* that lack the two *E. coli pyk* genes ^29^. Proteins were purified using ammonium sulfate fractionation and extensive dialysis ^5,9,27^. Because this is a partial purification, accurate estimates of hLPYK concentrations were not possible and therefore *k*_*cat*_ values could not be determined. Enzymatic activity was measured in a 96-well plate format using a lactate dehydrogenase/NADH-coupled assay to monitor the change in A_340_ at different substrate PEP concentrations. Due to the reliance of our assay on monitoring enzyme activity to first evaluate PEP affinity, no other parameters could be evaluated for variants that lacked catalytic activity. For all active variants, the PEP concentration that was equal to ½ maximal activity was designated as *K*_*app-PEP*_. This *K*_*app-PEP*_ value was simultaneously determined over a concentration range of each allosteric effector (Ala and Fru-1,6-BP). For each ligand, the *K*_*app-PEP*_ response to effector concentration was fit to:

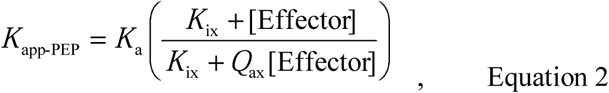

where *K*_*a*_ is the apparent affinity for ligand A (*e.g*., substrate *K*_*a-PEP*_) in the absence of ligand X (*e.g*., Ala or Fru-1,6-BP); and *K*_*ix*_ is the affinity for ligand X (*e.g*., *K*_*ix-Ala*_ and *K*_*ix-FBP*_) in the absence of ligand A (Table I). In Equation 2, *Q*_*ax*_ is the allosteric coupling constant defined by ^30-33^ as:

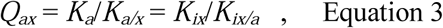

where *K*_*a/x*_ is the apparent affinity for ligand A in the presence of saturating ligand X; and *K*_*ix/a*_ is the affinity of the protein for ligand X in the presence of saturating ligand A. Other parameters are as defined for Equation 2.

In this study, each set of mutant proteins with substitutions at the same position were assayed concurrently. Each set also included a wildtype sample, which provided nine wildtype replicates. These wildtype replicates were used to estimate the error range for each wildtype parameter (Supplemental Table I).

### Classifying substitution patterns

Once the five functional parameters were determined for each variant, data were analyzed using the RheoScale calculator ^2^. This program uses all variants available for each position to perform several histogram analyses. Two of the three resulting RheoScale scores are useful to describe positions that make substantial functional contributions and have been well-described ^2^. In contrast, the current work to identify neutral positions demanded rigorous understanding of the RheoScale “neutral” score. This score is derived from the fraction of variants for which a functional parameter falls in the same histogram bin as that for the wildtype protein (*e.g*., all variants with *K*_*app-PEP*_ values equivalent to that of wildtype). To carry out these analyses, the RheoScale calculator has built-in recommendations for bin size and bin number. However, all histogram analyses require an empirical assessment of the most appropriate binning strategy. Since this was the first study in which we identified positions with strong neutral scores, it was appropriate to further develop the statistical considerations used to generate this score.

In the first iteration of RheoScale^2^, the experimental error was one of several considerations used to determine bin size. In our prior studies, other factors were more critical to the bin size. However, for neutrality, the error must be the dominant consideration: Substitutions at a position should have functional parameter measurements that, within error, are equivalent to wildtype protein. Therefore, we used the nine wildtype replicates of hLPYK to determine the width of the wildtype bin (Supplemental Table 2). To isolate the effects of amino acid substitution from variation in the other reagents used for enzyme assays, all variant data for each position were analyzed using a wildtype bin centered on the wildtype value collected on the same day. We found this hybrid use of wildtype data in the histogram analyses (*i.e*., the error derived from the average of all wildtype replicates, the bin centered on the wildtype data from the same day) to be the most conservative approach for identifying neutral substitutions and, subsequently, for assigning neutral positions. The alternative use of the nine-replicate wildtype average would give a higher false detection rate of neutral positions.

We further considered substitutions with values that fell just outside of the wildtype bin. Due to the expectation that wildtype replicates constitute a Gaussian distribution, some variant values that fell just outside of the wildtype bin might occupy the tails of the distribution. Thus, we considered adding weighted “sub-bins” immediately adjacent to each side of the wildtype bin. We chose to weight them with weighting scores less than 1 because there is some probability that these “near-wildtype” values indeed differ from wildtype and therefore should contribute less to the neutral score. The sub-bins were 1/5^th^ the size of the wildtype bin with weights ranging from 0.5 to 0.1. However, these additions had little influence on the final neutral score (Supplemental Figure 6). Therefore, these modifications were not included in the reported neutral scores. Nonetheless, the exercise to consider neighboring sub-bins added confidence that the bin size determined by wildtype error was optimal to evaluate neutral scores for evaluated positions.

## Results

hLPYK has many “functions.” We have developed a semi-high throughput assay that can simultaneously characterize substrate affinity (*K*_*app-PEP*_ for the substrate, phosphoenolpyruvate), binding of the allosteric inhibitor binding (*K*_*ix-Ala*_ for alanine), binding of the allosteric activator (*K*_*ix-FBP*_ for fructose-1,6-bisphosphate), allosteric coupling between the PEP and fructose-1,6-bisphosphate binding events (*Q*_*ax-FBP*_), and allosteric coupling between the PEP and Ala binding events (*Q*_*ax-Ala*_). By having a method that reports on many functions at once, we have found hLPYK to be a useful model to evaluate individual amino acid positions based on functional outcomes due to substitution^2,3^. In particular, our past studies have identified substitutions at nonconserved positions in and near the allosteric binding sites that modulated multiple functional parameters^2,3,5-9^ (Figure 1, green dots).

As a next question, we are interested in identifing the characteristics that determine when a nonconserved position does vs. does not influence function. The latter group of positions are termed “neutral” positions and can tolerate most, if not all, amino acid substitutions with no influence on the function of the protein. To address that next question, we wish to compare the biochemical, biophysical and structural characteristics of functionally neutral and important nonconserved positions. Thus, our goal in the current study was to identify a comparison set of neutral positions in hLPYK.

For the initial identification of potential neutral positions to be experimentally tested, we first considered three metrics commonly assumed to be associated with neutral positions. However, existing experimental data for hLPYK showed that the sensitivities of all three metrics were insufficient to guide experimental studies. (i) Extreme nonconservation, which can be quantified as sequence entropy (Equation 1), is often interpreted to indicate that the position lacks evolutionary constraints and thus is likely to be neutral. However, when we compared our previous experimental work^2,3^ to sequence entropy scores, positions across the full range could influence function (Figure 1). (ii) Surface exposed amino acid residues have fewer structural constraints than buried residues and are often assumed to tolerate a wide range of substitutions. However, in hLPYK, some positions in the allosteric sites are among the “surface exposed” residues (Figure 2), and therefore, surface exposure, in isolation, is not sufficient to predict a neutral position. Furthermore, a whole protein alanine scanning-mutation study of hLPYK^9^ identified many surface positions – far from the binding and active sites – that had substantial effects on function (Figure 2). (iii) Alanine scanning-mutagenesis are often used in isolation to identify functionally important protein positions. However, in our study of hLPYK^9^, several positions could be mutated to alanine with little functional consequence. Nonetheless, function was modified when those same positions were mutated to other residues^2,3^. In summary, neither the functional nonconserved positions nor the neutral positions could be uniquely identified using sequence entropy, surface exposure, or outcomes from alanine-scanning mutagenesis.

**Figure 2.**
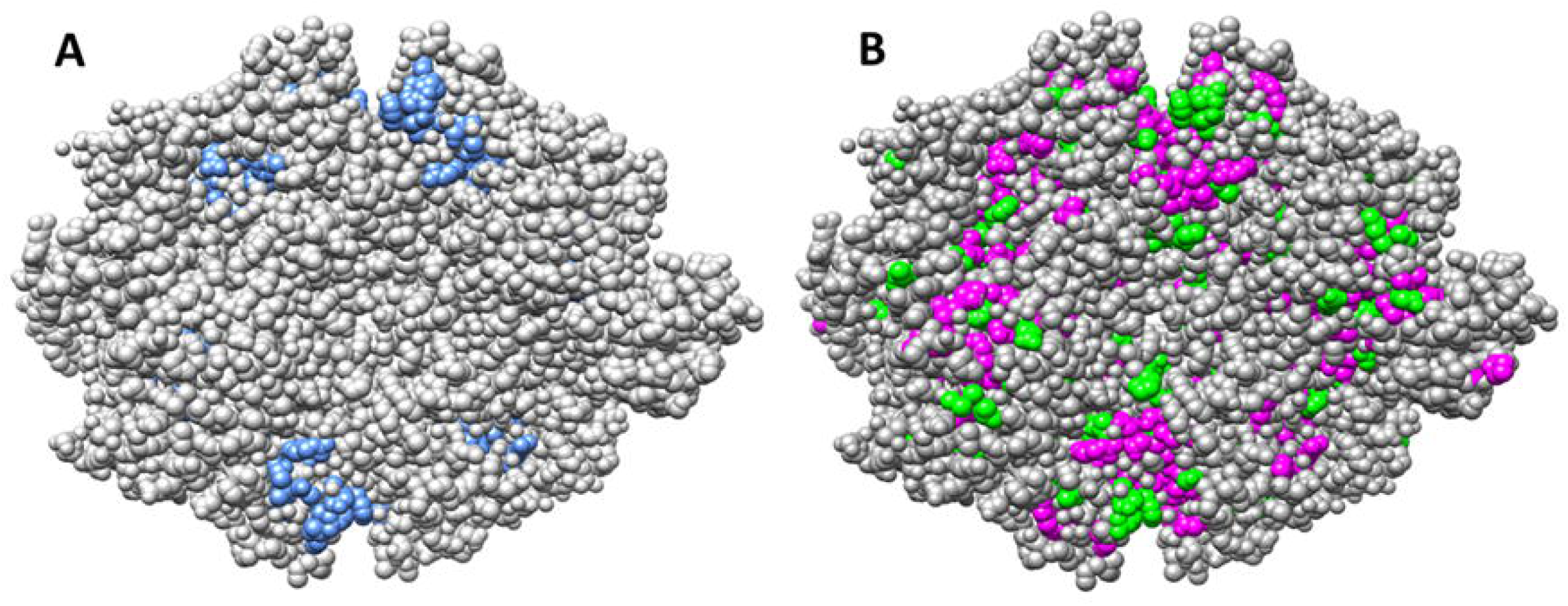
Surface exposure does not identify functionally neutral positions in the hLPYK homotetramer. Panel A shows the surface exposure of residues located in the two allosteric binding sites and previously evaluated for functional roles ^2,3,5,27^. Panel B highlights surface exposed positions at which substitution to alanine ^9^ causes either greater than a ten-fold change in one of 5 functional parameters (magenta) or between five- and ten-fold change in one of those five parameters (green). A tetramer of hLPYK from the PDB:4IMA structure was used to create this figure ^8^.

Thus, we next considered what other approaches might be useful to identify functionally neutral positions in hLPYK. We reasoned that sequence entropy only quantifies the extent of change observed at each position but does *not* distinguish whether the change occurs as part of an evolutionary pattern. Indeed, dozens of sequence analyses have been developed to identify patterns of change within a sequence alignment. As described in Methods, some nonconserved positions show a pattern of amino acid changes that mirrors the overall phylogeny of the protein family; experiments confirm that such “phylogenetic” positions play important functional roles ^14-16, 34, 35^. In other cases, pairs of nonconserved positions show correlated amino acid changes that are often associated with experimentally determined functional or structural roles ^17,18,20-25^.

For this work, we reasoned that positions exhibiting any pattern of evolutionary change should be excluded when attempting to identify neutral positions. Therefore, we combined several methods of classifying nonconserved positions into functional subclasses to score the likelihood that nonconserved positions exhibited a distinct pattern of change during evolution: Rather than noting the positions with the highest manifestation of a pattern, we identified the positions that were *least* likely to exhibit each of these patterns of change. From these, we generated a composite score to identify the positions that were overall *least likely* to exhibit any pattern in its amino acid changes (“least-patterned” positions). Notably, the correlation between the least patterned scores and sequence entropy scores was only −0.42 (Spearman nonparametric coefficient), which indicates that the least patterned score had significant differences from sequence entropy. Furthermore, alanine substitutions available for some of the 20 least patterned PYK positions all showed less than 4-fold change relative to wildtype (Figure 3).

**Figure 3.**
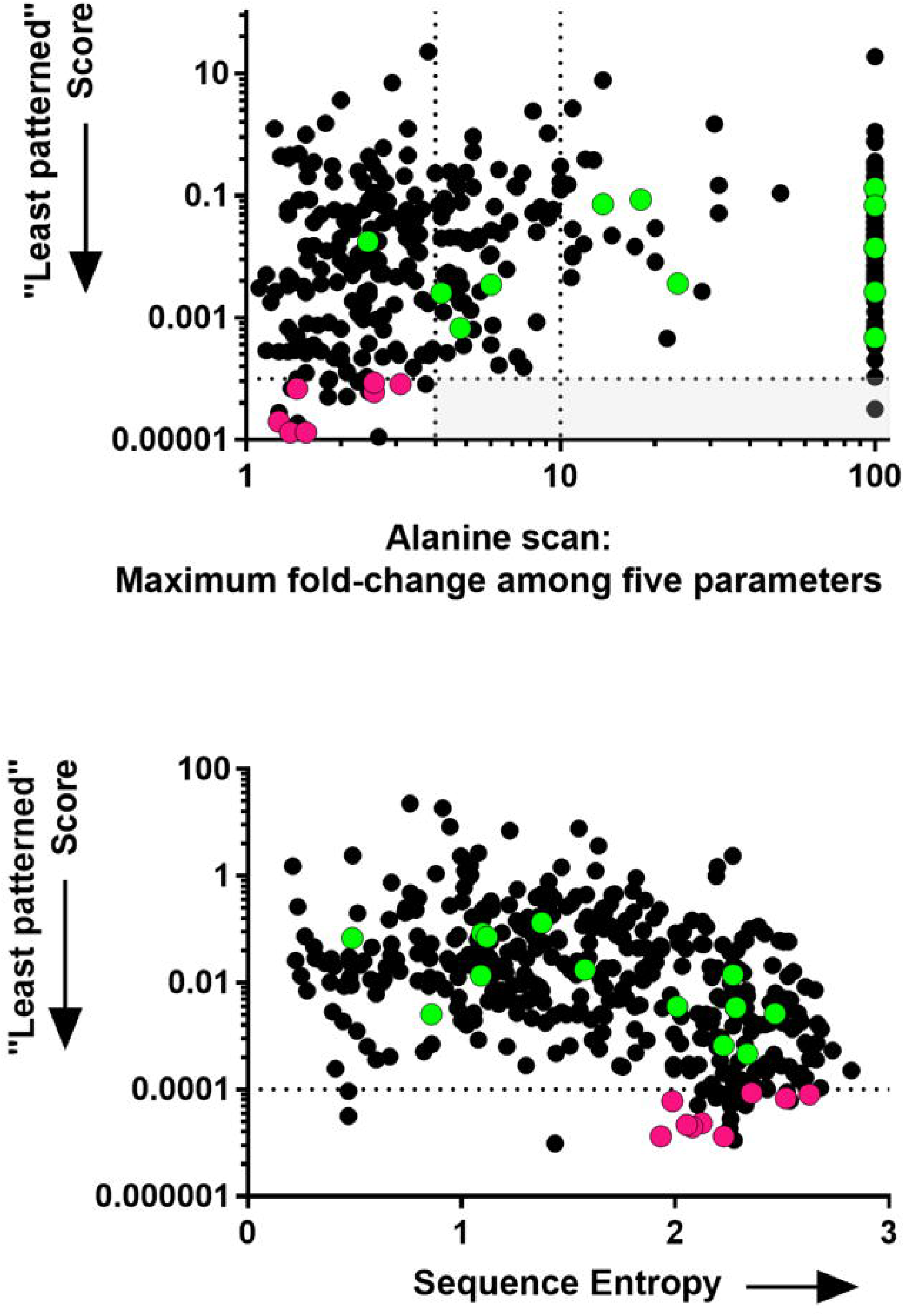
“Least patterned” scores calculated for hLPYK. Least patterned scores (black dots) were calculated for positions in the PYK sequence alignment, as described in the text. Smaller values indicate that the position shows more random changes in the sequence alignment. Co-evolutionary, and therefore least patterned, scores cannot be calculated for highly conserved positions (because they do not change); therefore, highly conserved positions are not represented in either panel. The overlaid green dots show the positions in the allosteric sites ^2,3,5,27^ and the magenta dots show the positions substituted in this study. (Top) “Least patterned” scores versus fold-change from the alanine-scanning mutagenesis study ^9^. Dotted lines and the light gray box are used to illustrate that all but one of the least patterned positions has < 4-fold change; the second dotted line at fold-change = 10 corresponds to the dotted line in Figure 1. (Bottom) “Least patterned” scores versus sequence entropies show that these two metrics have different rank orders for the PYK amino acid positions.

For the twenty-two “least patterned” hLPYK positions, we next assessed all 19 substitutions at each position using a machine learning algorithm– SNAP– that was specifically designed to predict neutral substitutions^26^. Of the top twenty-two positions, 20 passed the SNAP test. This list included positions: 75, 101, 138, 147, 151, 196, 199, 206, 208, 210, 214, 239, 246, 268, 381, 388, 412, 470, 511, and 517.

Before beginning experiments, we next verified that no previously documented substitutions alter function by cross-checking against existing experimental data. The first data set was the ∼260 missense polymorphisms in human erythrocyte PYK isozyme (hRPYK) associated with non-spherocytic hemolytic anemia^4,36^. Most mutations in hRPYK are also present in hLPYK, because the same gene encodes the two proteins; the gene encoding hLPYK has an alternative start site used for the expression of the longer erythrocyte hRPYK isozyme ^37^. The second large data set that we considered included the positions important to allosteric regulation that were identified *via* a whole-protein, alanine-scanning mutagenesis study of 431 positions in hLPYK ^9^. A final data set included 17 positions in the hLPYK allosteric sites that, when substituted, modulated allosteric regulation to varying degrees ^2,3,5,27^. Of the potentially neutral positions identified by the “least patterned” scores, these existing experimental data disqualified position 511 from further consideration (Supplemental Figure 3). A disease-associated mutation has been reported at position 246^38^; however, as discussed in the supplement, we believe this is an erroneous assignment. Thus, our study included position 246 among the positions chosen for experimental verification.

Of the remaining list, we noted that positions 196, 199, 206, 208, 210, and 214 clustered on the primary sequence and we chose to experimentally evaluate 199, 206, 208, 210, and 214. Positions 75, 138, 246, and 412 were selected to represent positions scattered throughout the rest of the primary sequence. At least nine substitutions at each position (Figure 4), for a total of 117 variant proteins, were analyzed using >22,000 activity assays to assess substitution effects on ligand affinity/binding and allosteric coupling. Quantitative values for each parameter (Table I) were determined for each variant protein (Supplemental Table 3). For each parameter, the data for all variants at each position were then evaluated using the RheoScale calculator, as described in Materials and Methods, to calculate “neutral” scores.

**Figure 4.**
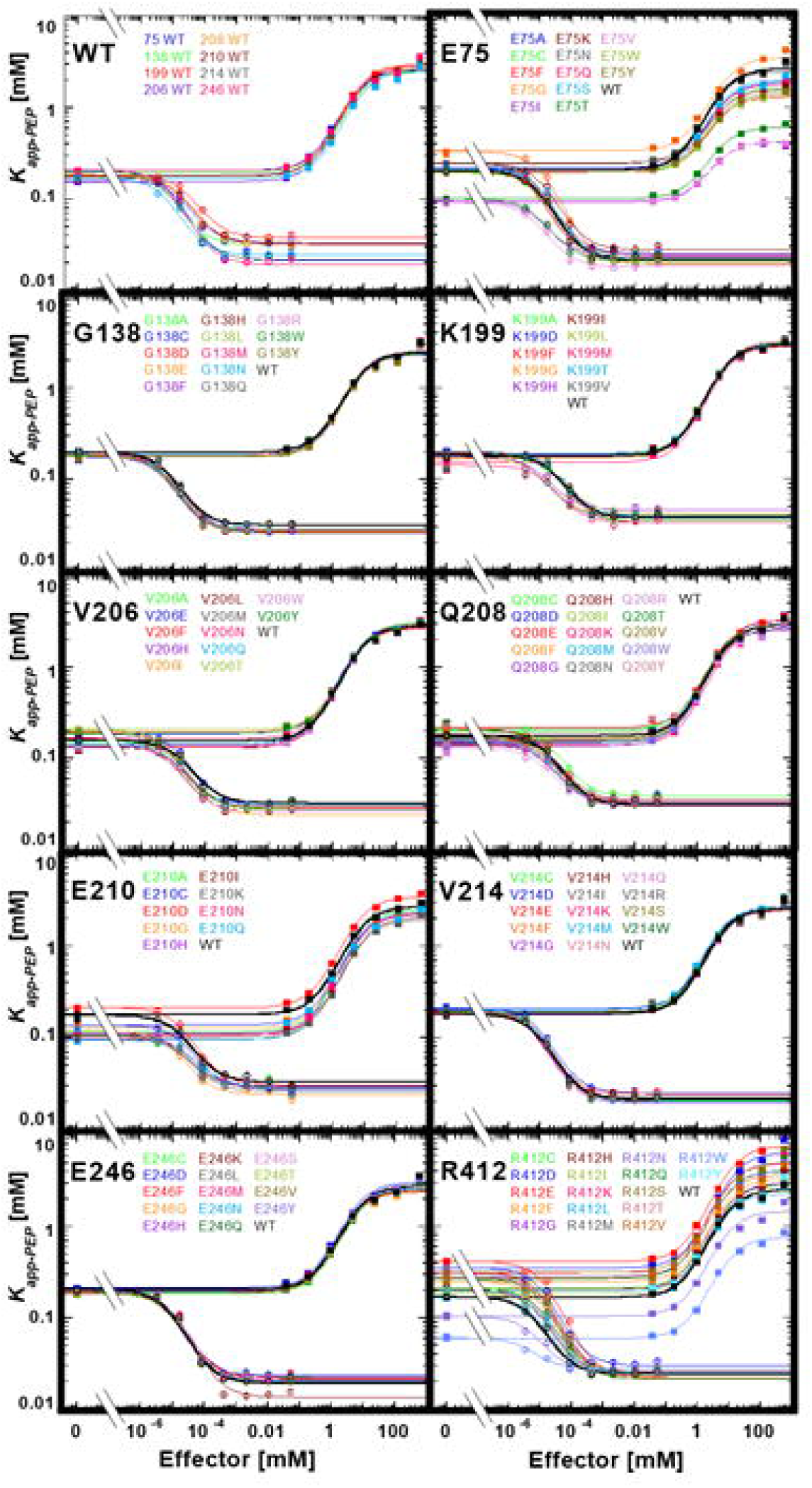
Functional characterization of hLPYK variants. The response of the apparent affinity of PEP (determined as the concentration of PEP that results in ½ maximal velocity) was measured as a function of effector concentrations for all hLPYK variants. Values for *K*_*app-PEP*_ at various concentrations of both the allosteric inhibitor Ala and the allosteric activator Fru-1,6-BP were determined ^5, 9, 27^. In this figure, all variants made at one position are shown on the individual panels. Lines are best fits to Equation 2, and all fitted parameters for all variants are listed in Supplemental Table 3. A key for viewing how fit values are represented in this figure is included in Supplemental Figure 2. Error bars (most often smaller than the data symbols) represent errors of the fit for the *K*_*app-PEP*_ values. The assays for all variants at a given position were carried out on the same day, along with a wildtype protein that is included in each panel. To represent day-to-day variability, all wildtype data sets are shown in the first panel. As described in Materials and Methods, the range of parameters from these wildtype replicates was used to establish bin size in RheoScale calculations.

The definition of neutrality can be customized to various levels of restrictiveness. A highly restrictive definition would require a position to be insensitive to all 19 possible substitutions. A less restrictive definition would exclude proline and glycine, both of which alter the polypeptide backbone, from the list of substitutions expected to be accommodated at one position. Additionally, SNAP predicted several positions in which tryptophan was the only detrimental substitution, which could be due to its large size. Of the three exceptions considered, the proline assumption was supported by our experimental results: Of seven positions with proline substitutions, only positions 138 and 412 retained any enzymatic activity. Therefore, data for proline substitutions were included in data tables in the supplement but were not included in our assessments of mutational outcomes. In contrast, glycine and tryptophan were accommodated in several positions without altering function. Therefore, glycine and tryptophan substitutions were included in the RheoScale calculation of neutral scores.

A simplification required to make the project tractable was to avoid generating all possible substitutions at each tested position. Instead, we relied on a random sampling of substitutions. Even with the choices to select only nine positions in hLPYK and work with fewer than 20 substitutions, this study generated 117 variant proteins that were evaluated with >22,000 assays. Consistent with our original goal, when outcomes were summarized, substitutions at positions 138, 214, and 246 had little influence on *any* of the parameters evaluated. This neutrality is reflected in the perfect RheoScale neutral scores of 1 for all parameters (Figure 5).

**Figure 5.**
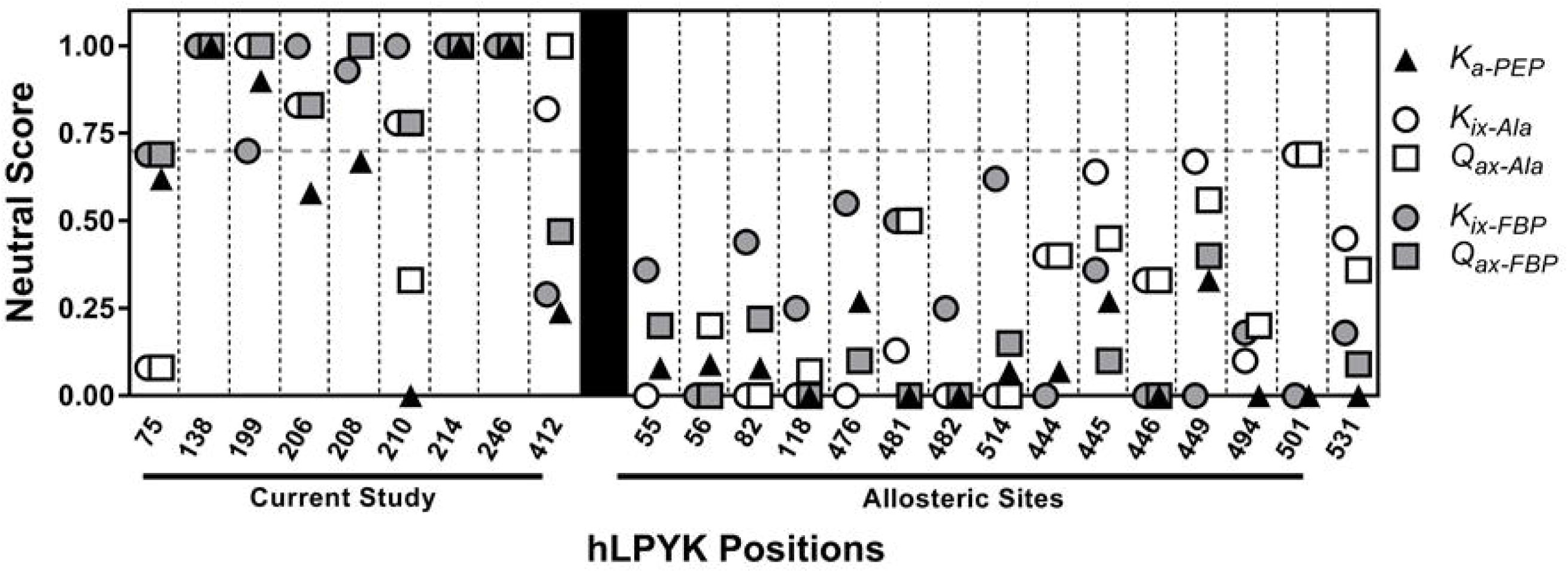
Neutral scores for positions in hLPYK. Left: Parameters derived from data in Figure 4 were used in the RheoScale calculator to quantify the neutral character of each position. hLPYK positions are noted on the x-axis. The effects of substitutions on PEP affinity (*K*_*a-PEP*_), Ala binding (*K*_*ix-Ala*_), PEP/Ala allosteric coupling (*Q*_*ax-Ala*_), Fru-1,6-BP binding (*K*_*ix-FBP*_), and PEP/Fru-1,6-BP allosteric coupling (*Q*_*ax-FBP*_) were used to calculate neutral scores for each parameter. Dashed vertical lines separate the scores for individual positions. Right: Neutral scores for positions in allosteric sites evaluated elsewhere^3^ are included here to offer a contrast of positions that exhibit a lower range of neutral scores.

In addition, we considered that less-perfect RheoScale neutral scores could still reflect a high tendency for neutrality (*i.e*., if non-neutral substitutions caused only modest effects and none of those non-neutral substitutions caused a severe outcome on function). We examined the primary data in Figure 4 and the analysis of that data in Figure 5 and Supplemental Table 3. We also compared outcomes from this study with results from two previous hLPYK studies ^2,3^ that targeted sites with known functional importance. Empirically, neutral scores above 0.70 distinguished the positions of this study from those positions known to have large functional changes upon substitution. This comparison was used to derive a significance threshold of 0.70 for strong neutral scores (Figure 5). Using that threshold, we concluded that positions 199, 206, and 208, although not perfectly neutral, are near-neutral.

We acknowledge that we can never test all possible functions for a protein. Furthermore, as we reviewed in a previous discussion about the impacts of substituting nonconserved positions^13^, some substitutions might alter protein stability. However, within the context of the specific functions evaluated in this study, our approach to identifying neutral positions successfully enriched for positions for which substitutions have neutral or modest outcomes on substrate and ligand binding or on allosteric regulation.

## Discussion

In order to identify which characteristics of nonconserved positions dictate when a position does or does not contribute to function, a comparison set of neutral positions is critical. Thus, the initial goal of the current project was to identify neutral positions in hLPYK. However, hLPYK has 543 positions to consider, and examination of previous experiments ruled out three commonly assumed approaches to selecting neutral positions: High sequence entropy (Figures 1), surface exposure (Figure 2), and identification *via* an alanine-scanning mutation study. Thus, to define a tractable set of positions for experimental verification, we were first required to develop a strategy to identify positions most likely to be neutral in hLPYK.

The strategy we used was to combine several existing sequence analyses (phylogenetic and coevolutionary) to generate a set of “least patterned” positions that was further assessed with a substitution predictor (SNAP) for neutrality. For the nine positions tested experimentally, three positions (138, 214, and 246) were neutral for all five functional parameters (Figure 5). A fourth position, 199, also had high neutral scores and, along with two more positions (206 and 208), met the 0.7 threshold. These were classified as “near-neutral.” Furthermore, substitutions at the remaining positions in this study generally had small to modest effects on function, with fewer substitutions that altered function in comparison to positions located in functionally-important binding sites (Figure 5; Supplemental Figure 4). Thus, we did indeed identify functionally neutral positions based on biochemical assays.

Overall, the group of “neutral” positions selected by our approach was enriched for a higher percent of neutral substitutions, as reflected in their larger neutrality scores, as compared to positions located near allosteric binding sites (Supplemental Figure 5). Thus, our approach for selecting potentially-neutral positions might be generally useful for guiding experiments to identify neutral positions and substitutions in other proteins. However, we note that we have not tested this method for false negatives (*i.e*., neutral positions that are not predicted to be neutral by our strategy), nor did we experimentally test all 19 substitutions for each position. We also note that, although the current work included SNAP in the selection pipeline ^26,39,40^, we did not assess whether SNAP alone (or the updated version of the algorithm released after our analyses^39^) would have been sufficient to enrich for neutral and near-neutral positions. Finally, we note that large numbers of analyses have been designed to identify patterns of evolutionary change in sequence alignments; it is possible that some combination of pattern scores might not identify neutral positions as successfully as in this study. Future studies should consider the relative weights of different analyses to the least patterned scores.

Another consideration is whether the least patterned scores are protein family-dependent. This would first require a consideration of whether the absolute magnitudes of least patterned scores have meaning, which in turn, requires understanding whether the raw scores from the contributing analyses are comparable among protein families. However, aside from sequence entropy, we are unaware of any such comparisons. For example, co-evolution scores are commonly used to identify positions that most strongly co-evolve within a family, but we are not aware of studies to compare whether there is any significance to one family having stronger scoring pairs of positions than a second family.

A second consideration for generalizing neutral positions identification requires considering the boundaries of sequence diversity chosen to define the protein family. We previously considered the effect of family size on sequence analyses ^13,18,41,42^; different sequence identity cutoffs delineate superfamilies, families, and subfamilies. The work described in the current work utilized the largest sequence space that could be identified with PYK functionality (99% to <15% sequence identity). Future sequence-based analyses of neutral positions should systematically vary the boundaries of the protein family to determine their contributions to the “least patterned” signal.

Indeed, the question of the sequence family boundaries raises a question about the extensibility of current neutral positions to other homologs in the PYK family. Given the large sequence alignment used, the expectation might be that the neutral and near-neutral positions identified in hLPYK will be universally neutral (or near-neutral) in the entire PYK family. Another possibility is that some of the near-neutral positions in hLPYK are neutral in other PYK isozymes. Alternatively, we previously showed that the locations of evolutionarily-constrained positions can differ among subfamilies^18^. That is, the whole protein family had a common set of strongly conserved and co-evolving positions, and each subfamily had additional strongly conserved and co-evolving positions in unique locations (*i.e*., different columns of the sequence alignment) for each subfamily. In corollary, each PYK subfamily may also have a set of unique neutral positions, in addition to a family-wide set of neutral and near-neutral positions.

In an interesting parallel to the current study, Bromberg and colleagues published the algorithm “fuNTRp” during the final stages of our current work. This algorithm is designed to predict whether a given position will exhibit neutral, rheostat, or toggle substitution behavior^43^. Their approach employed machine learning and was trained and validated on substitution outcomes derived from deep mutational scanning (“DMS”). In DMS, a library of protein variants is expressed in a cellular context; cells compete for several generations; high throughput DNA sequencing is used to determine the prevalence of each substitution; substitution prevalence is used to infer altered protein function^44,45^. As such, DMS intrinsically includes a biological threshold for neutrality, but that threshold can change with altered selection conditions^46^. In contrast, our current work utilized biochemical characterization that directly monitors functional change of the protein. To our knowledge, the ability of fuNTRp to predict biochemically neutral positions has not yet been tested.

Even if our strategy for identifying neutral positions lacks general applicability in other proteins, the current identification of neutral positions has immediate value to benchmark and/or train new prediction algorithms. Likely because of the historic focus on conserved positions^1^, Bromberg *et al*. noted that the development of predictive strategies was inhibited by insufficient numbers of experimentally-confirmed (or reported) neutral substitutions^40^. A variety of neutral substitutions and positions have been predicted computationally (*e.g*.,^47^), but many need to be validated with biochemical assays. More examples of neutral substitutions and positions might be gleaned from the ever-growing database of “deep mutational scanning” studies (*e.g*., ^44,45,48^). However, these results are a composite of functional change, “solubility” changes^49^, altered interactions with chaperones, and are sensitive to the biological conditions used for selection^46^. Likewise, “non-pathogenic” substitutions extracted from disease and exome databases (*e.g*., ^47^) may also be context-dependent. Therefore, the biochemically neutral positions of this study provide a rare dataset.

Another interesting note is that the strategy we used to select the 20 possibly-neutral positions did *not* consider their locations on the protein structure. Therefore, we were surprised at the structural patterns that were apparent when the predicted positions were mapped onto the structure of hLPYK^8^. A cluster of predicted neutral positions (196, 199, 206, 208, 210, and 214) resides on the B-domain (Figure 6). The second group of predicted neutral positions showed a pattern when mapped onto the A-domain (Figure 6). The A-domain has a classic TIM barrel fold with eight parallel β-strands surrounded by eight α-helices. The predicted neutral positions mapped to the centers of solvent-exposed α-helices in the TIM barrel. Interestingly, this complements work from another study of other TIM barrel proteins ^2,50^ in which two classes of functionally important positions showed structural clustering at the top and bottom of β-sheets in the TIM barrel. It will be interesting to determine whether these are general patterns for TIM barrels and/or if patterns of position types are common in other secondary folds.

**Figure 6.**
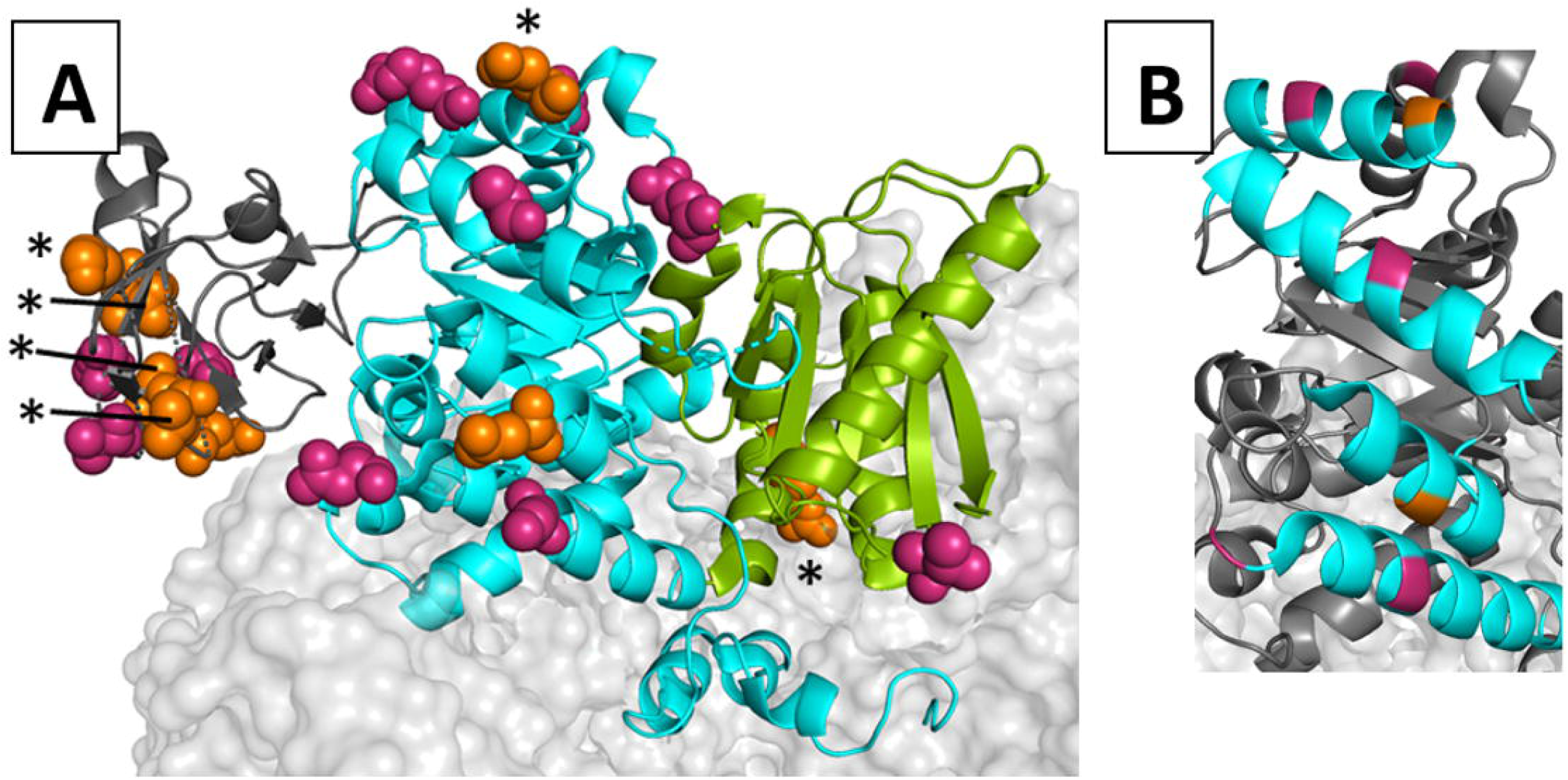
Locations of potentially-neutral positions on the hLPYK homotetramer. The structure used for this model was PDB:4IMA ^8^. One monomer is shown as a ribbon, whereas neighboring subunits were rendered with a light gray surface. A) On the highlighted monomer, its A-, B-, and C-domains are colored cyan, dark gray, and green, respectively. The N-terminal domain that is often differentiated in other works is not distinguished from the A domain in this color scheme. The side chains of positions predicted to be neutral positions are shown with orange (experimentally tested herein) and magenta (proposed but not tested) spacefilling. Because position 138 is not ordered in this structure, position 137 was highlighted as a proxy. Neutral and near-neutral positions are marked with an asterisk. B) Several potentially-neutral positions (orange and magenta) are located in the middle of helices (cyan) of the TIM barrel of the A-domain. This pattern occurs even though the TIM barrel in hLPYK is not a continuous chain. Instead, the B-domain is inserted in the middle of the sequence that forms the TIM barrel of the A-domain.

Overall, this study successfully identified neutral positions in hLPYK. Those positions can be used in future comparative studies to determine what structure/function properties determine when a nonconserved position does *vs*. does not influence functions. In future studies, we envision structural studies to evaluate how these two types of nonconserved positions accommodate substitutions, as well as evaluations of protein dynamics (potentially using hydrogen/deuterium exchange mass spectrometry^51^) to determine if the intrinsic dynamic properties of the regions where the two types of nonconserved positions reside dictate contributions to function. However, independent of what techniques are used in the future, this initial identification of neutral positions will facilitate those future comparative studies.

## Acknowledgments

This research was supported by NIH grants GM115340 and GM118589 and a grant from the W.M. Keck Foundation.

## References

1. Gray VE, Kukurba KR, Kumar S. Performance of computational tools in evaluating the functional impact of laboratory-induced amino acid mutations. Bioinformatics. 2012;28(16):2093–6.

2. Hodges AM, Fenton AW, Dougherty LL, et al. RheoScale: A tool to aggregate and quantify experimentally determined substitution outcomes for multiple variants at individual protein positions. Hum Mutat. 2018;39(12):1814–26.

3. Wu T, Swint-Kruse L, Fenton AW. Functional tunability from a distance: Rheostat positions influence allosteric coupling between two distant binding sites. Sci Rep. 2019;9(1):16957.

4. Pendergrass DC, Williams R, Blair JB, et al. Mining for allosteric information: Natural mutations and positional sequence conservation in pyruvate kinase. IUBMB Life. 2006;58(1):31–8.

5. Ishwar A, Tang Q, Fenton AW. Distinguishing the interactions in the fructose 1,6-bisphosphate binding site of human liver pyruvate kinase that contribute to allostery. Biochemistry. 2015;54(7):1516–24.

6. Fenton AW, Tang Q. An activating interaction between the unphosphorylated n-terminus of human liver pyruvate kinase and the main body of the protein is interrupted by phosphorylation. Biochemistry. 2009;48(18):3816–8.

7. McFarlane JS, Ronnebaum TA, Meneely KM, et al. Changes in the allosteric site of human liver pyruvate kinase upon activator binding include the breakage of an intersubunit cation-pi bond. Acta Crystallogr F Struct Biol Commun. 2019;75(Pt 6):461–69.

8. Holyoak T, Zhang B, Deng J, et al. Energetic coupling between an oxidizable cysteine and the phosphorylatable N-terminus of human liver pyruvate kinase. Biochemistry. 2013;52(3):466–76.

9. Tang Q, Fenton AW. Whole-protein alanine-scanning mutagenesis of allostery: a large percentage of a protein can contribute to mechanism. Hum Mutat. 2017;38(9):1132–43.

10. Blair JB. Regulatory Properties of Hepatic Pyruvate Kinase. In: Veneziale CM, ed., The Regulation of Carbohydrate Formation and Utilization in Mammals. Baltimore: University Park Press, 1980.

11. Hall ER, Cottam GL. Isozymes of pyruvate kinase in vertebrates: their physical, chemical, kinetic and immunological properties. Int J Biochem. 1978;9(11):785–93.

12. Hall TA. BioEdit: a user-friendly biological sequence alignment editor and analysis program for Windows. Nucleic Acids Symposium Series. 1999;41(95-98.

13. Swint-Kruse L. Using Evolution to Guide Protein Engineering: The Devil IS in the Details. Biophys J. 2016;111(1):10–8.

14. Ye K, Vriend G, Ap IJ. Tracing evolutionary pressure. Bioinformatics. 2008;24(7):908–15.

15. Mihalek I, Res I, Lichtarge O. Evolutionary trace report_maker: a new type of service for comparative analysis of proteins. Bioinformatics. 2006;22(13):1656–7.

16. Gu X, Zou Y, Su Z, et al. An update of DIVERGE software for functional divergence analysis of protein family. Mol Biol Evol. 2013;30(7):1713–9.

17. Brown CA, Brown KS. Validation of coevolving residue algorithms via pipeline sensitivity analysis: ELSC and OMES and ZNMI, oh my! PLoS One. 2010;5(6):e10779.

18. Parente DJ, Swint-Kruse L. Multiple co-evolutionary networks are supported by the common tertiary scaffold of the LacI/GalR proteins. PLoS One. 2013;8(12):e84398.

19. Parente DJ, Ray JC, Swint-Kruse L. Amino acid positions subject to multiple coevolutionary constraints can be robustly identified by their eigenvector network centrality scores. Proteins. 2015;83(12):2293–306.

20. Gobel U, Sander C, Schneider R, et al. Correlated mutations and residue contacts in proteins. Proteins. 1994;18(4):309–17.

21. Olmea O, Rost B, Valencia A. Effective use of sequence correlation and conservation in fold recognition. J Mol Biol. 1999;293(5):1221–39.

22. Olmea O, Valencia A. Improving contact predictions by the combination of correlated mutations and other sources of sequence information. Fold Des. 1997;2(3):S25–32.

23. Fodor AA, Aldrich RW. Influence of conservation on calculations of amino acid covariance in multiple sequence alignments. Proteins. 2004;56(2):211–21.

24. Kass I, Horovitz A. Mapping pathways of allosteric communication in GroEL by analysis of correlated mutations. Proteins. 2002;48(4):611–7.

25. Dekker JP, Fodor A, Aldrich RW, et al. A perturbation-based method for calculating explicit likelihood of evolutionary co-variance in multiple sequence alignments. Bioinformatics. 2004;20(10):1565–72.

26. Bromberg Y, Yachdav G, Rost B. SNAP predicts effect of mutations on protein function. Bioinformatics. 2008;24(20):2397–8.

27. Tang Q, Alontaga AY, Holyoak T, et al. Exploring the limits of the usefulness of mutagenesis in studies of allosteric mechanisms. Hum Mutat. 2017.

28. Valentini G, Chiarelli LR, Fortin R, et al. Structure and function of human erythrocyte pyruvate kinase. Molecular basis of nonspherocytic hemolytic anemia. J Biol Chem. 2002;277(26):23807–14.

29. Fenton AW, Hutchinson M. The pH dependence of the allosteric response of human liver pyruvate kinase to fructose-1,6-bisphosphate, ATP, and alanine. Arch Biochem Biophys. 2009;484(16-23.

30. Reinhart GD. The determination of thermodynamic allosteric parameters of an enzyme undergoing steady-state turnover. Arch Biochem Biophys. 1983;224(1):389–401.

31. Reinhart GD. Linked-function origins of cooperativity in a symmetrical dimer. Biophys Chem. 1988;30(2):159–72.

32. Reinhart GD. Quantitative analysis and interpretation of allosteric behavior. Methods Enzymol. 2004;380(187-203.

33. Weber G. Ligand binding and internal equilibria in proteins. Biochemistry. 1972;11(5):864–78.

34. Lichtarge O, Bourne HR, Cohen FE. An evolutionary trace method defines binding surfaces common to protein families. J Mol Biol. 1996;257(2):342–58.

35. Gu X, Vander Velden K. DIVERGE: phylogeny-based analysis for functional-structural divergence of a protein family. Bioinformatics. 2002;18(3):500–1.

36. Canu G, De Bonis M, Minucci A, et al. Red blood cell PK deficiency: An update of PK-LR gene mutation database. Blood Cells Mol Dis. 2016;57(100-9.

37. Noguchi T, Yamada K, Inoue H, et al. The L- and R-type isozymes of rat pyruvate kinase are produced from a single gene by use of different promoters. J Biol Chem. 1987;262(29):14366–71.

38. Berghout J, Higgins S, Loucoubar C, et al. Genetic diversity in human erythrocyte pyruvate kinase. Genes Immun. 2012;13(1):98–102.

39. Hecht M, Bromberg Y, Rost B. Better prediction of functional effects for sequence variants. BMC Genomics. 2015;16 Suppl 8(S1.

40. Bromberg Y, Rost B. SNAP: predict effect of non-synonymous polymorphisms on function. Nucleic Acids Res. 2007;35(11):3823–35.

41. Tungtur S, Parente DJ, Swint-Kruse L. Functionally important positions can comprise the majority of a protein’s architecture. Proteins. 2011;79(5):1589–608.

42. Meinhardt S, Manley MW, Jr., Parente DJ, et al. Rheostats and toggle switches for modulating protein function. PLoS One. 2013;8(12):e83502.

43. Miller M, Vitale D, Kahn PC, et al. funtrp: identifying protein positions for variation driven functional tuning. Nucleic Acids Res. 2019;47(21):e142.

44. Fowler DM, Fields S. Deep mutational scanning: a new style of protein science. Nat Methods. 2014;11(8):801–7.

45. Roscoe BP, Thayer KM, Zeldovich KB, et al. Analyses of the effects of all ubiquitin point mutants on yeast growth rate. J Mol Biol. 2013;425(8):1363–77.

46. Mavor D, Barlow K, Thompson S, et al. Determination of ubiquitin fitness landscapes under different chemical stresses in a classroom setting. Elife. 2016;5(

47. Valiaho J, Faisal I, Ortutay C, et al. Characterization of all possible single-nucleotide change caused amino acid substitutions in the kinase domain of Bruton tyrosine kinase. Hum Mutat. 2015;36(6):638–47.

48. Gray VE, Hause RJ, Fowler DM. Analysis of Large-Scale Mutagenesis Data To Assess the Impact of Single Amino Acid Substitutions. Genetics. 2017;207(1):53–61.

49. Klesmith JR, Bacik JP, Wrenbeck EE, et al. Trade-offs between enzyme fitness and solubility illuminated by deep mutational scanning. Proc Natl Acad Sci U S A. 2017;114(9):2265–70.

50. Chan YH, Venev SV, Zeldovich KB, et al. Correlation of fitness landscapes from three orthologous TIM barrels originates from sequence and structure constraints. Nat Commun. 2017;8(14614.

51. Prasannan CB, Villar MT, Artigues A, et al. Identification of regions of rabbit muscle pyruvate kinase important for allosteric regulation by phenylalanine, detected by H/D exchange mass spectrometry. Biochemistry. 2013;52(11):1998–2006.

