## Supplement for "Identification of biochemically neutral positions in liver pyruvate kinase"

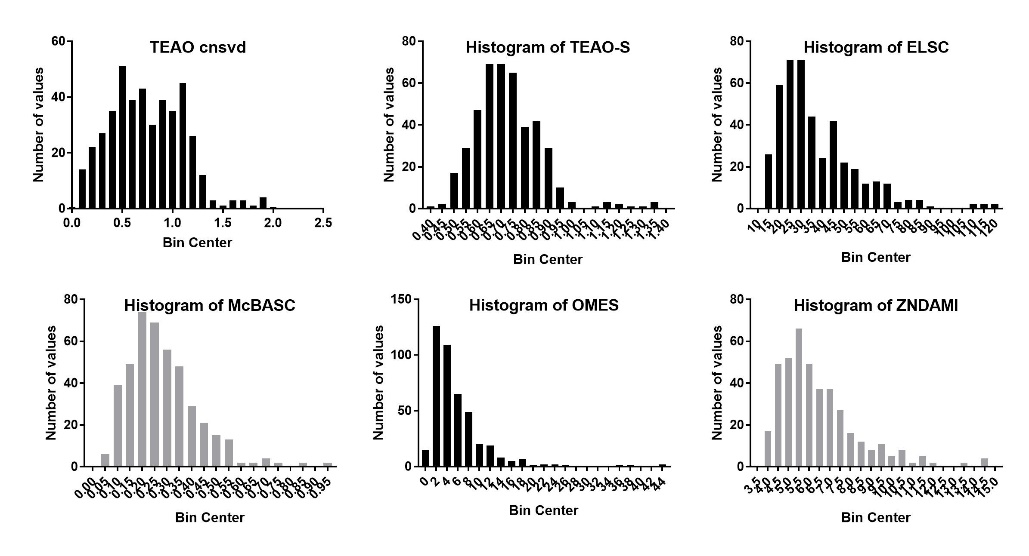

**Supplemental Figure 1.** Score distributions for the six sequence analyses used in combination to generate the least patterned score. All are all some form of a Gaussian curve with different skew and kurtosis. Note that the least patterned scoring uses the “not significant” end of every distribution (low values in co-evolution, high score values in TEAO).

**Supplemental Definitions: Additional RheoScale Scores**

In our earlier studies, we identified three classes of amino acid positions based on their substitution outcomes ^2, 3^. (i) As explored in this manuscript, a perfectly neutral position accepts any amino acid side chain without altering functional parameters. (ii) “Toggle” positions are those for which most substitutions are catastrophic ^2, 3^. (iii) “Rheostat” positions are those for which a range of amino acid substitutions result in a wide range of functional outcomes, from wildtype (or better), to various intermediate levels, to dead. In addition to the neutral score discussed in this manuscript, for a set of substitutions at a given position, the RheoScale calculator ^2^ also reports a score for the fraction of variants that abolish function (“toggle” score) and a score that reflects the prevalence of intermediate outcomes (“rheostat” score). These terms and scores are included in the Tables and Figures of this supplement.

**Supplemental Results: The dependency of histogram analysis on the error of each parameter**

The role of the error associated with each parameter on determining bin size is emphasized by considering *K_a-PEP_* and *Q_ax-FBP_* for position 208 (Figures 4 and 5). The wildtype range/bin size for *K_a-PEP_* is small, resulting in a small wildtype bin size. Therefore, the visible variation in *K_a-PEP_* values resulting from substitutions at position 208 results in a neutral score for *K_a-PEP_* that is below 0.7. In contrast to *K_a-PEP_*, the wildtype range/bin size for *Q_ax-FBP_* is large. Upon visual inspection of the wildtype replicates and recalling the ratio definition for *Q_ax_* in Equation 3, the increased error associated with *Q_ax-FBP_* appears to be a result of increased variance in the *K_a/x-PEP/FBP_* value (lower plateau at high Fru-1,6-BP concentration; Figure 4).

In contrast, in the example of position 208, there are more differences in *K_a-PEP_* values (plateau at low Fru-1,6-BP concentrations; Figure 4) than *K_a/x-PEP/FBP_* values. Nonetheless, the resulting *Q_ax-FBP_* values for 208 variants fall into the greater wildtype *Q_ax-FBP_* bin size, even though fewer of the *K_a-PEP_* values for 208 fell into the smaller wildtype *K_a-PEP_* bin size*.* As a result of this dependence on neutral score calculation on the wildtype error for each parameter, the visible variability in *K_a-PEP_* for 208 variants results in a low neutral *K_a-PEP_* score, but a high *Q_ax-FBP_* neutral score.

**Supplemental Discussion: E246K as a cause of disease**

Our data indicates neutrality for position246. After experiments were begun, an updated pyruvate kinase deficiency disease database ^36^ included the E246K-equivalent in the list of substitutions in erythrocyte PYK that cause disease. (The actual entry is E277K due to the 31 additional N-terminal residues in RPYK that are absent in LPYK). The E246K-equivalent was identified in a population screen: individuals screened were not reported to have pyruvate kinase deficiency ^38^. Furthermore, in contrast to a statement in the original manuscript, no previous publications report the E277K (i.e., the E246K-equivalent) mutation in RPYK as a cause of pyruvate kinase deficiency. Therefore, there is no available data to challenge our assignment of position 246 as neutral.

**Supplemental Figure 2.** A visual aid to help the reader identify parameters in Figure 4. Binding constants (K_ix-Ala_ and K_ix-FBP_) are derived from the inflection points.

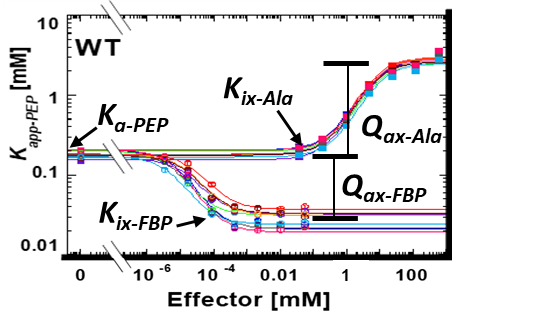

**Supplemental Table 1: Wildtype averages**. The error from wildtype replicates was used to establish bin size for neutral score determination by the RheoScale calculator. Because the data from Wu *et al*. is included as a comparison in tables, the wildtype averages from that study are included here for comparison.

|  | **Current study** | | **Previous study** ^3^ | |
| --- | --- | --- | --- | --- |
|  | **Average**  **(replicates)** | **Standard deviation of independent determinations** | **Average**  **(replicates)** | **Standard deviation of independent determinations** |
| ***K_a-PEP_* (mM)** | 0.19  (n=18) | 0.02 | 0.24  (n=10) | 0.02 |
| ***K_ix-ala_* (mM)** | 0.43  (n=9) | 0.06 | 0.31  (n=5) | 0.04 |
| ***K_ix-FBP_* (µM)** | 0.09  (n=9) | 0.03 | 0.19  (n=5) | 0.08 |
| ***Q_ax-ala_*** | 0.068  (n=9) | 0.009 | 0.073  (n=5) | 0.009 |
| ***Q_ax-FBP_*** | 7  (n=9) | 2 | 14  (n=5) | 3 |

**Supplemental Table 2: Histogram parameters used in the RheoScale calculator for each functional parameter.** The default bin number is the minimum of the following: 1) median number of variants per position, 2) mean number of variants per variant, and 3) the range divided by error. The RheoScale default bin size was calculated by splitting a specified range of data by the recommended (default) number of bins; therefore, bin size was sometimes but not always determined by error. In contrast, as we noted, the primary importance of error on considerations of neutrality, the neutral scores were calculated by the number of variants that fell inside a bin centered on the wildtype average and that was sufficiently wide to capture all observed wildtype values.

|  | **Neutral** | | |
| --- | --- | --- | --- |
|  | **RheoScale default bin number (ranging from WT to dead)** | **RheoScale default bin size (log) used for rheostat and toggle score** **determina­tions** ^2, 3^ | **WT capture bin size (log) used for neutral score determinations in this work** |
| ***K_a-PEP_*** | 14 | 0.219 | 0.146 |
| ***K_ix-Ala_*** | 14 | 0.330 | 0.175 |
| ***K_ix-FBP_*** | 14 | 0.381 | 0.639 |
| ***Q_ax-Ala_*** | 10 | 0.137 | 0.186 |
| ***Q_ax-FBP_*** | 3 | 0.561 | 0.372 |

**Supplemental Table 3: Experimental parameters determined for each of the variants in this study**. The parameters were determined from the data in Figure 4 using Equation 2. Reported errors are errors of the fits. The values in the table below were used to calculate the neutral scores of Figure 5 and the rheostat and toggle scores of Supplemental Figure 4.

| **Protein** | ***K_a-PEP_* [mM]** | ***K_ix-Ala_* [mM]** | ***Q_ax-Ala_*** | ***K_ix-FBP_* [µM]** | ***Q_ax-FBP_*** |
| --- | --- | --- | --- | --- | --- |
| Glu75 | | | | | |
| Wildtype (E75) | 0.205±0.003 | 0.41±0.02 | 0.078±0.002 | 0.090±0.006 | 9.4±0.3 |
| E75A | 0.219±0.001 | 0.62±0.02 | 0.12±0.01 | 0.090±0.007 | 8.7±0.2 |
| E75C | 0.195±0.003 | 0.64±0.03 | 0.12±0.01 | 0.073±0.007 | 7.9±0.3 |
| E75F | 0.207±0.004 | 0.78±0.07 | 0.17±0.01 | 0.062±0.006 | 9.0±0.3 |
| E75G | 0.332±0.003 | 0.63±0.03 | 0.096±0.002 | 0.23±0.01 | 16.9±0.6 |
| E75I | 0.094±0.002 | 1.0±0.1 | 0.22±0.01 | 0.035±0.004 | 4.0±0.1 |
| E75K | 0.246±0.003 | 0.80±0.04 | 0.19±0.01 | 0.11±0.01 | 8.9±0.3 |
| E75N | 0.243±0.004 | 0.50±0.03 | 0.099±0.003 | 0.093±0.008 | 10.6±0.3 |
| E75Q | 0.200±0.002 | 0.55±0.03 | 0.11±0.01 | 0.098±0.006 | 8.3±0.2 |
| E75S | 0.210±0.003 | 0.53±0.02 | 0.10±0.01 | 0.11±0.01 | 9.7±0.3 |
| E75T | 0.101±0.002 | 0.78±0.06 | 0.16±0.01 | 0.042±0.006 | 4.3±0.2 |
| E75V | 0.090±0.002 | 1.0±0.1 | 0.21±0.01 | 0.034±0.005 | 5.0±0.2 |
| E75W | 0.194±0.003 | 0.77±0.04 | 0.14±0.01 | 0.089±0.008 | 8.9±0.3 |
| E75Y | 0.202±0.004 | 0.76±0.05 | 0.14±0.01 | 0.07±0.01 | 10.0±0.4 |
| Gly138 | | | | | |
| Wildtype (G138) | 0.200±0.003 | 0.53±0.02 | 0.081±0.002 | 0.041±0.004 | 6.3±0.2 |
| G138A | 0.186±0.003 | 0.50±0.03 | 0.077±0.003 | 0.043±0.005 | 6.7±0.2 |
| G138C | 0.178±0.005 | 0.50±0.04 | 0.077±0.004 | 0.034±0.005 | 6.7±0.4 |
| G138D | 0.191±0.003 | 0.56±0.03 | 0.075±0.002 | 0.036±0.004 | 7.2±0.2 |
| G138E | 0.187±0.005 | 0.57±0.05 | 0.077±0.004 | 0.036±0.004 | 6.6±0.3 |
| G138F | 0.182±0.004 | 0.57±0.04 | 0.076±0.003 | 0.040±0.005 | 6.4±0.2 |
| G138H | 0.189±0.004 | 0.62±0.04 | 0.075±0.003 | 0.053±0.007 | 7.1±0.3 |
| G138L | 0.183±0.004 | 0.55±0.04 | 0.077±0.003 | 0.033±0.004 | 6.7±0.3 |
| G138M | 0.199±0.003 | 0.52±0.02 | 0.079±0.002 | 0.038±0.005 | 7.1±0.2 |
| G138N | 0.192±0.003 | 0.54±0.03 | 0.079±0.002 | 0.035±0.005 | 7.1±0.3 |
| G138Q | 0.188±0.004 | 0.55±0.04 | 0.075±0.003 | 0.042±0.005 | 7.3±0.3 |
| G138R | 0.189±0.004 | 0.51±0.03 | 0.080±0.004 | 0.042±0.005 | 6.8±0.3 |
| G138W | 0.2003±0.006 | 0.53±0.05 | 0.077±0.005 | 0.036±0.005 | 7.0±0.3 |
| G138Y | 0.178±0.006 | 0.51±0.05 | 0.075±0.004 | 0.039±0.006 | 6. 7±0.4 |
| Lys199 | | | | | |
| Wildtype (K199) | 0.183±0.004 | 0.37±0.02 | 0.060±0.002 | 0.14±0.02 | 4.9±0.2 |
| K199A | 0.190±0.003 | 0.44±0.02 | 0.061±0.001 | 0.13±0.02 | 5.4±0.3 |
| K199D | 0.195±0.002 | 0.43±0.02 | 0.064±0.001 | 0.16±0.02 | 5.1±0.2 |
| K199F | 0.192±0.002 | 0.42±0.01 | 0.063±0.001 | 0.12±0.02 | 5.1±0.2 |
| K199G | 0.169±0.003 | 0.36±0.02 | 0.059±0.002 | 0.04±0.01 | 4.1±0.1 |
| K199H | 0.180±0.004 | 0.43±0.03 | 0.060±0.002 | 0.040±0.006 | 3.8±0.1 |
| K199I | 0.196±0.003 | 0.40±0.02 | 0.060±0.002 | 0.14±0.02 | 5.9±0.2 |
| K199L | 0.188±0.002 | 0.37±0.02 | 0.064±0.001 | 0.14±0.01 | 4.4±0.1 |
| K199M | 0.148±0.003 | 0.36±0.02 | 0.054±0.002 | 0.07±0.01 | 4.3±0.2 |
| K199T | 0.195±0.003 | 0.40±0.02 | 0.062±0.002 | 0.108±0.009 | 4.9±0.2 |
| K199V | 0.182±0.004 | 0.43±0.03 | 0.062±0.002 | 0.040±0.005 | 4.6±0.2 |
| Val206 | | | | | |
| Wildtype (V206) | 0.158±0.003 | 0.36±0.02 | 0.055±0.001 | 0.09±0.01 | 4.9±0.2 |
| V206A | 0.156±0.003 | 0.33±0.02 | 0.053±0.001 | 0.069±0.008 | 5.5±0.2 |
| V206E | 0.190±0.004 | 0.41±0.02 | 0.065±0.002 | 0.10±0.02 | 6.1±0.3 |
| V206F | 0.136±0.002 | 0.31±0.02 | 0.051±0.002 | 0.057±0.006 | 4.6±0.1 |
| V206H | 0.140±0.002 | 0.33±0.02 | 0.051±0.001 | 0.058±0.006 | 4.5±0.1 |
| V206I | 0.198±0.006 | 0.44±0.02 | 0.074±0.002 | 0.07±0.02 | 8.1±0.6 |
| V206L | 0.133±0.003 | 0.30±0.02 | 0.047±0.002 | 0.056±0.006 | 5.2±0.2 |
| V206M | 0.143±0.002 | 0.32±0.01 | 0.050±0.001 | 0.055±0.008 | 4.5±0.2 |
| V206N | 0.167±0.003 | 0.36±0.02 | 0.057±0.002 | 0.064±0.009 | 5.8±0.2 |
| V206Q | 0.147±0.003 | 0.31±0.02 | 0.051±0.001 | 0.049±0.005 | 4.7±0.1 |
| V206T | 0.208±0.006 | 0.53±0.03 | 0.072±0.002 | 0.07±0.02 | 7.5±0.5 |
| V206W | 0.136±0.002 | 0.31±0.02 | 0.048±0.001 | 0.050±0.005 | 5.3±0.2 |
| V206Y | 0.189±0.004 | 0.46±0.03 | 0.062±0.002 | 0.062±0.009 | 6.6±0.3 |
| Gln208 | | | | | |
| Wildtype (Q208) | 0.180±0.003 | 0.41±0.02 | 0.061±0.002 | 0.11±0.02 | 5.7±0.2 |
| Q208C | 0.203±0.005 | 0.44±0.02 | 0.067±0.002 | 0.13±0.03 | 5.3±0.3 |
| Q208D | 0.173±0.003 | 0.37±0.01 | 0.055±0.001 | 0.08±0.01 | 4.9±0.2 |
| Q208E | 0.216±0.005 | 0.49±0.03 | 0.066±0.002 | 0.09±0.02 | 6.2±0.3 |
| Q208F | 0.160±0.003 | 0.36±0.02 | 0.059±0.002 | 0.08±0.01 | 4.9±0.2 |
| Q208G | 0.153±0.002 | 0.35±0.02 | 0.056±0.001 | 0.08±0.01 | 5.1±0.2 |
| Q208H | 0.163±0.002 | 0.39±0.01 | 0.059±0.001 | 0.12±0.01 | 4.7±0.2 |
| Q208I | 0.166±0.002 | 0.37±0.01 | 0.056±0.001 | 0.09±0.01 | 5.3±0.2 |
| Q208K | 0.142±0.002 | 0.42±0.02 | 0.053±0.001 | 0.063±0.006 | 4.3±0.1 |
| Q208M | 0.156±0.003 | 0.36±0.02 | 0.052±0.001 | 0.070±0.008 | 4.8±0.2 |
| Q208N | 0.148±0.003 | 0.37±0.02 | 0.057±0.002 | 0.07±0.01 | 4.4±0.2 |
| Q208R | 0.136±0.002 | 0.37±0.02 | 0.055±0.001 | 0.045±0.006 | 3.8±0.2 |
| Q208T | 0.209±0.005 | 0.42±0.02 | 0.072±0.003 | 0.09±0.01 | 6.6±0.3 |
| Q208V | 0.173±0.003 | 0.39±0.02 | 0.058±0.001 | 0.09±0.01 | 5.7±0.2 |
| Q208W | 0.140±0.003 | 0.31±0.01 | 0.052±0.001 | 0.070±0.009 | 4.3±0.2 |
| Q208Y | 0.178±0.005 | 0.45±0.03 | 0.065±0.003 | 0.07±0.01 | 4.8±0.2 |
| Glu210 | | | | | |
| Wildtype (E210) | 0.178±0.004 | 0.41±0.02 | 0.067±0.002 | 0.10±0.01 | 5.3±0.2 |
| E210A | 0.112±0.002 | 0.35±0.02 | 0.047±0.001 | 0.08±0.01 | 3.8±0.1 |
| E210C | 0.138±0.002 | 0.38±0.02 | 0.059±0.002 | 0.12±0.02 | 5.0±0.2 |
| E210D | 0.215±0.003 | 0.38±0.02 | 0.063±0.002 | 0.14±0.02 | 7.0±0.3 |
| E210G | 0.123±0.003 | 0.37±0.02 | 0.054±0.002 | 0.057±0.007 | 4.9±0.2 |
| E210H | 0.111±0.002 | 0.34±0.01 | 0.051±0.001 | 0.08±0.01 | 3.7±0.1 |
| E210I | 0.107±0.001 | 0.40±0.01 | 0.053±0.001 | 0.066±0.007 | 3.6±0.1 |
| E210K | 0.099±0.001 | 0.38±0.02 | 0.052±0.001 | 0.080±0.009 | 3.72±0.08 |
| E210N | 0.102±0.001 | 0.32±0.01 | 0.047±0.001 | 0.11±0.01 | 3.33±0.07 |
| E210Q | 0.096±0.001 | 0.28±0.01 | 0.041±0.001 | 0.11±0.01 | 3.24±0.08 |
| Val214 | | | | | |
| Wildtype (V214) | 0.184±0.002 | 0.43±0.02 | 0.071±0.002 | 0.077±0.006 | 8.4±0.2 |
| V214C | 0.192±0.002 | 0.40±0.02 | 0.076±0.002 | 0.076±0.006 | 9.3±0.3 |
| V214D | 0.212±0.002 | 0.43±0.01 | 0.077±0.002 | 0.083±0.005 | 8.5±0.2 |
| V214E | 0.190±0.002 | 0.43±0.02 | 0.075±0.003 | 0.068±0.006 | 7.8±0.2 |
| V214F | 0.191±0.002 | 0.45±0.02 | 0.077±0.002 | 0.078±0.009 | 8.7±0.3 |
| V214G | 0.182±0.003 | 0.42±0.02 | 0.067±0.002 | 0.074±0.007 | 9.3±0.3 |
| V214H | 0.202±0.002 | 0.43±0.02 | 0.079±0.002 | 0.077±0.007 | 9.8±0.3 |
| V214I | 0.204±0.002 | 0.42±0.02 | 0.075±0.002 | 0.089±0.005 | 9.7±0.2 |
| V214K | 0.182±0.003 | 0.43±0.02 | 0.075±0.002 | 0.067±0.008 | 8.2±0.3 |
| V214M | 0.204±0.002 | 0.36±0.01 | 0.076±0.002 | 0.088±0.006 | 10.1±0.3 |
| V214N | 0.189±0.003 | 0.40±0.02 | 0.074±0.003 | 0.059±0.006 | 9.0±0.3 |
| V214Q | 0.194±0.003 | 0.45±0.02 | 0.078±0.003 | 0.071±0.005 | 8.7±0.2 |
| V214R | 0.187±0.003 | 0.42±0.02 | 0.073±0.002 | 0.074±0.008 | 8.6±0.4 |
| V214S | 0.189±0.002 | 0.40±0.02 | 0.074±0.002 | 0.074±0.006 | 8.8±0.3 |
| V214W | 0.189±0.002 | 0.40±0.02 | 0.073±0.002 | 0.075±0.007 | 8.8±0.3 |
| Glu246 | | | | | |
| Wildtype (E246) | 0.207±0.003 | 0.45±0.02 | 0.072±0.002 | 0.089±0.007 | 10.7±0.3 |
| E246C | 0.186±0.002 | 0.48±0.02 | 0.071±0.002 | 0.078±0.007 | 8.3±0.2 |
| E246D | 0.215±0.004 | 0.41±0.02 | 0.070±0.002 | 0.068±0.007 | 9.1±0.3 |
| E246F | 0.196±0.004 | 0.38±0.02 | 0.079±0.003 | 0.09±0.01 | 8.8±0.3 |
| E246G | 0.195±0.004 | 0.43±0.02 | 0.080±0.003 | 0.09±0.01 | 9.0±0.3 |
| E246H | 0.193±0.003 | 0.44±0.02 | 0.074±0.002 | 0.09±0.01 | 9.5±0.4 |
| E246K | 0.189±0.003 | 0.42±0.02 | 0.074±0.002 | 0.17±0.02 | 13.7±0.8 |
| E246L | 0.191±0.003 | 0.47±0.02 | 0.073±0.002 | 0.078±0.009 | 8.9±0.3 |
| E246M | 0.204±0.002 | 0.43±0.02 | 0.072±0.002 | 0.102±0.007 | 9.5±0.2 |
| E246N | 0.200±0.003 | 0.40±0.02 | 0.073±0.002 | 0.099±0.007 | 10.0±0.3 |
| E246Q | 0.197±0.002 | 0.40±0.01 | 0.078±0.002 | 0.089±0.008 | 9.9±0.3 |
| E246S | 0.197±0.002 | 0.43±0.02 | 0.075±0.002 | 0.098±0.008 | 9.6±0.3 |
| E246T | 0.190±0.003 | 0.44±0.02 | 0.067±0.002 | 0.093±0.007 | 9.8±0.3 |
| E246V | 0.189±0.002 | 0.38±0.02 | 0.067±0.002 | 0.09±0.01 | 9.9±0.4 |
| E246Y | 0.204±0.003 | 0.41±0.02 | 0.071±0.002 | 0.090±0.009 | 9.0±0.3 |
| Arg412 | | | | | |
| Wildtype (R412) | 0.170±0.002 | 0.52±0.02 | 0.067±0.002 | 0.043±0.006 | 7.0±0.2 |
| R412C | 0.212±0.003 | 0.54±0.02 | 0.074±0.002 | 0.085±0.009 | 8.7±0.3 |
| R412D | 0.353±0.003 | 0.65±0.02 | 0.055±0.001 | 0.140±0.009 | 11.8±0.3 |
| R412E | 0.415±0.004 | 0.56±0.02 | 0.055±0.002 | 0.28±0.02 | 18.6±0.7 |
| R412F | 0.248±0.003 | 0.61±0.02 | 0.075±0.002 | 0.101±0.008 | 9.5±0.2 |
| R412G | 0.203±0.002 | 0.66±0.02 | 0.067±0.001 | 0.070±0.007 | 8.1±0.3 |
| R412H | 0.205±0.002 | 0.52±0.02 | 0.070±0.002 | 0.091±0.007 | 7.9±0.2 |
| R412I | 0.185±0.002 | 0.51±0.02 | 0.075±0.002 | 0.09±0.01 | 7.7±0.3 |
| R412K | 0.311±0.004 | 0.51±0.02 | 0.064±0.001 | 0.14±0.01 | 12.9±0.4 |
| R412L | 0.168±0.002 | 0.54±0.03 | 0.074±0.002 | 0.052±0.005 | 6.6±0.2 |
| R412M | 0.195±0.003 | 0.59±0.03 | 0.067±0.002 | 0.069±0.008 | 8.1±0.3 |
| R412N | 0.104±0.002 | 0.57±0.03 | 0.071±0.003 | 0.024±0.003 | 4.1±0.1 |
| R412Q | 0.268±0.003 | 0.52±0.02 | 0.065±0.002 | 0.14±0.01 | 12.4±0.4 |
| R412S | 0.317±0.003 | 0.59±0.03 | 0.063±0.002 | 0.15±0.01 | 12.7±0.3 |
| R412T | 0.287±0.003 | 0.56±0.02 | 0.071±0.002 | 0.13±0.01 | 11.7±0.3 |
| R412V | 0.276±0.003 | 0.50±0.02 | 0.075±0.002 | 0.15±0.02 | 12.5±0.5 |
| R412W | 0.060±0.001 | 0.70±0.06 | 0.078±0.004 | 0.010±0.003 | 2.2±0.09 |
| R412Y | 0.199±0.002 | 0.62±0.02 | 0.076±0.002 | 0.092±0.007 | 8.1±0.2 |

In initial assays, proline substitutions abolished activity for positions 75, 199, 206, 208, and 210. No attempt to replicate these assays was attempted. Due to a general intolerance for proline, all proline data were removed from the analysis. However, proline substitutions were tolerated at positions 138 and 412. The data for those mutant proteins is as follows:

| **Protein** | ***K_a-PEP_* [mM]** | ***K_ix-Ala_* [mM]** | ***Q_ax-Ala_*** | ***K_ix-FBP_* [µM]** | ***Q_ax-FBP_*** |
| --- | --- | --- | --- | --- | --- |
| G138P | 0.167±0.003 | 0.53±0.03 | 0.073±0.002 | 0.036±0.005 | 6.1±0.2 |
| R412P | 1.32±0.01 | 1.33±0.05 | 0.132±0.002 | 1.5±0.1 | 48±2 |

**Supplemental Figure 3.** Data from the previous alanine-scanning mutagenesis study ^9^. Prior to the current experiments, the alanine-scanning data were used to pre-screen the top 20 “least-patterned” positions. The error bars shown are errors of the fit. For position 511 (red symbols), the alanine substitution altered multiple functional parameters relative to wildtype (black symbol and solid line). Therefore, we concluded that position 511 could not be perfectly neutral and did not further consider it in the current study. Note that we did not exclude the possibility that other substitutions at position 511 have neutral outcomes. Position 511 may, in fact, have intermediate neutral scores, like the other positions shown in Figure 5.

**Supplemental Figure 3.**

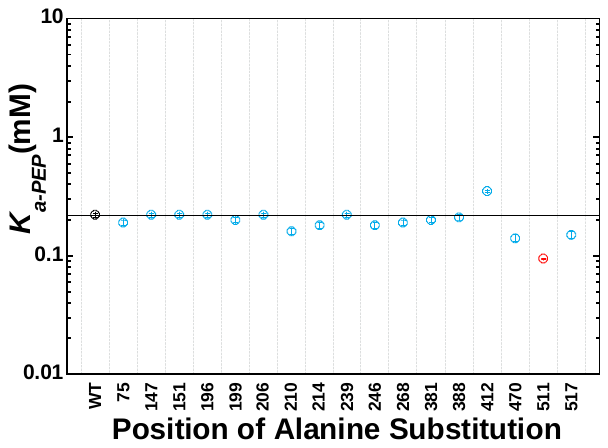

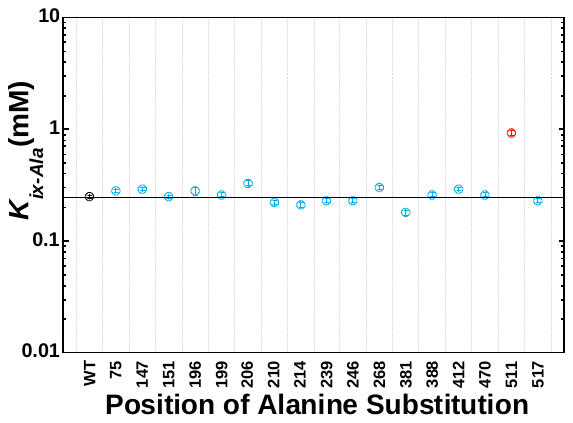

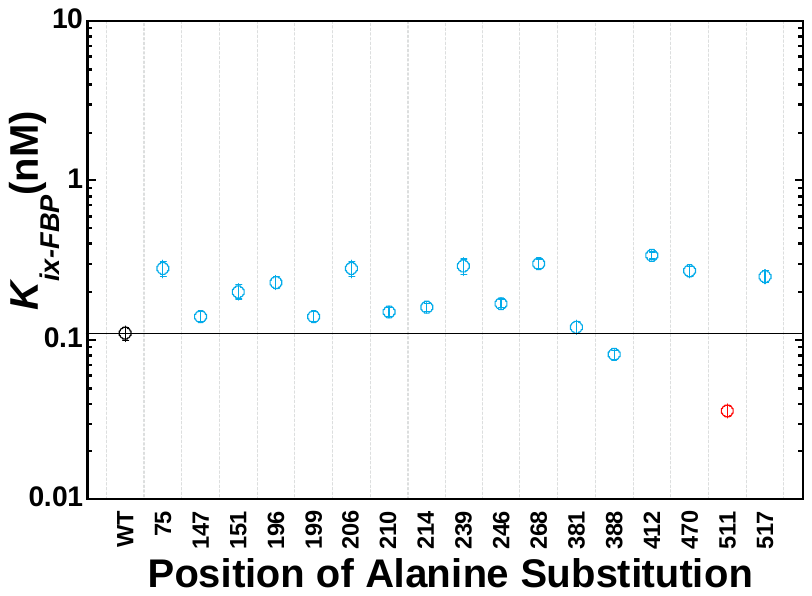

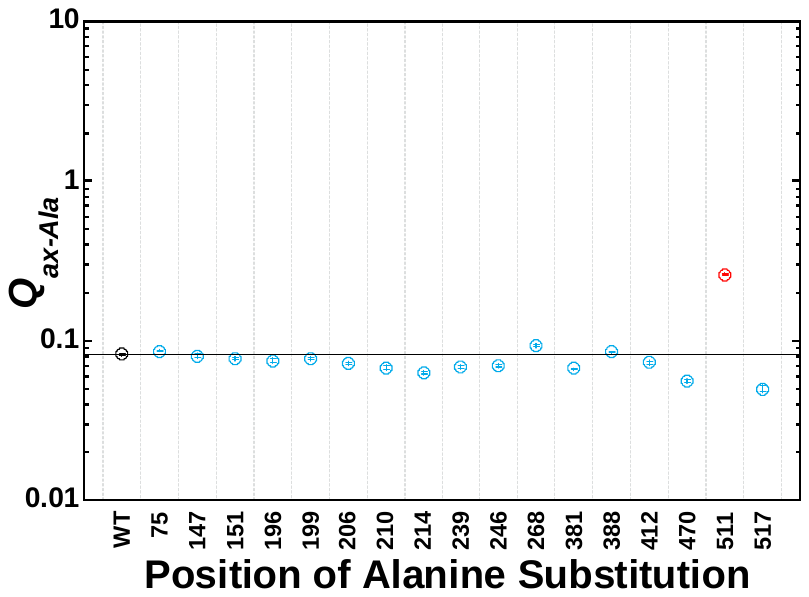

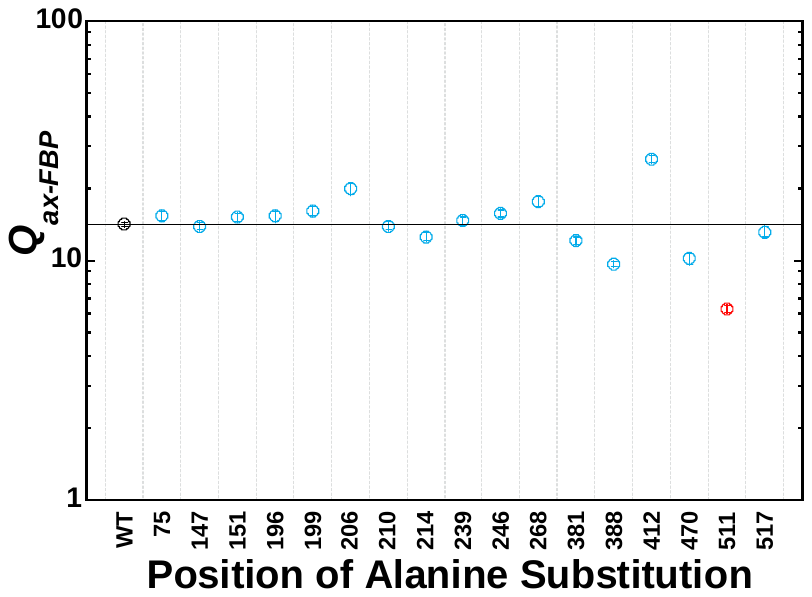

**Supplemental Figure 4. Comparisons of neutral, rheostat, and toggle RheoScale scores for positions of this study and positions located near allosteric binding sites of hLPYK.** The definitions of these terms as scores are described at the beginning of this Supplement. Ideally, the bin parameters used in histogram analyses would be identical for all datasets. This was possible for 4 of the 5 functional parameters. However, in the current dataset, the error determined from the wildtype *Q_ax-FBP_* values was larger than in the previous data set (Supplemental Tables 2 and 3). For the current *Q_ax-FBP_* data, RheoScale recommended a bin number of 3. That small of bin number was not meaningful for determining rheostat scores. Therefore, we did not calculate a rheostat score for *Q_ax-FBP_* for the positions evaluated in the current study.

In panel A, the dotted horizontal line at 0.7 was empirically determined to divide scores for positions with strong neutral character and scores of known functional positions. In panel B, the dotted line at 0.5 was previously determined^2^ as an indicator for strong rheostat character. In panel C, the dotted line at 0.64 for toggle scores was a previously determined^3^ as a cutoff for toggle characteristics. Note that for the positions of the current study, only one (position 75) has a significant rheostat score, and all toggle scores are zero. This suggests that the current strategy to identify these positions enriched results for neutral and modest substitutions, even though only a small fraction of positions have the potential to be perfectly neutral.

**Supplemental Figure 4**

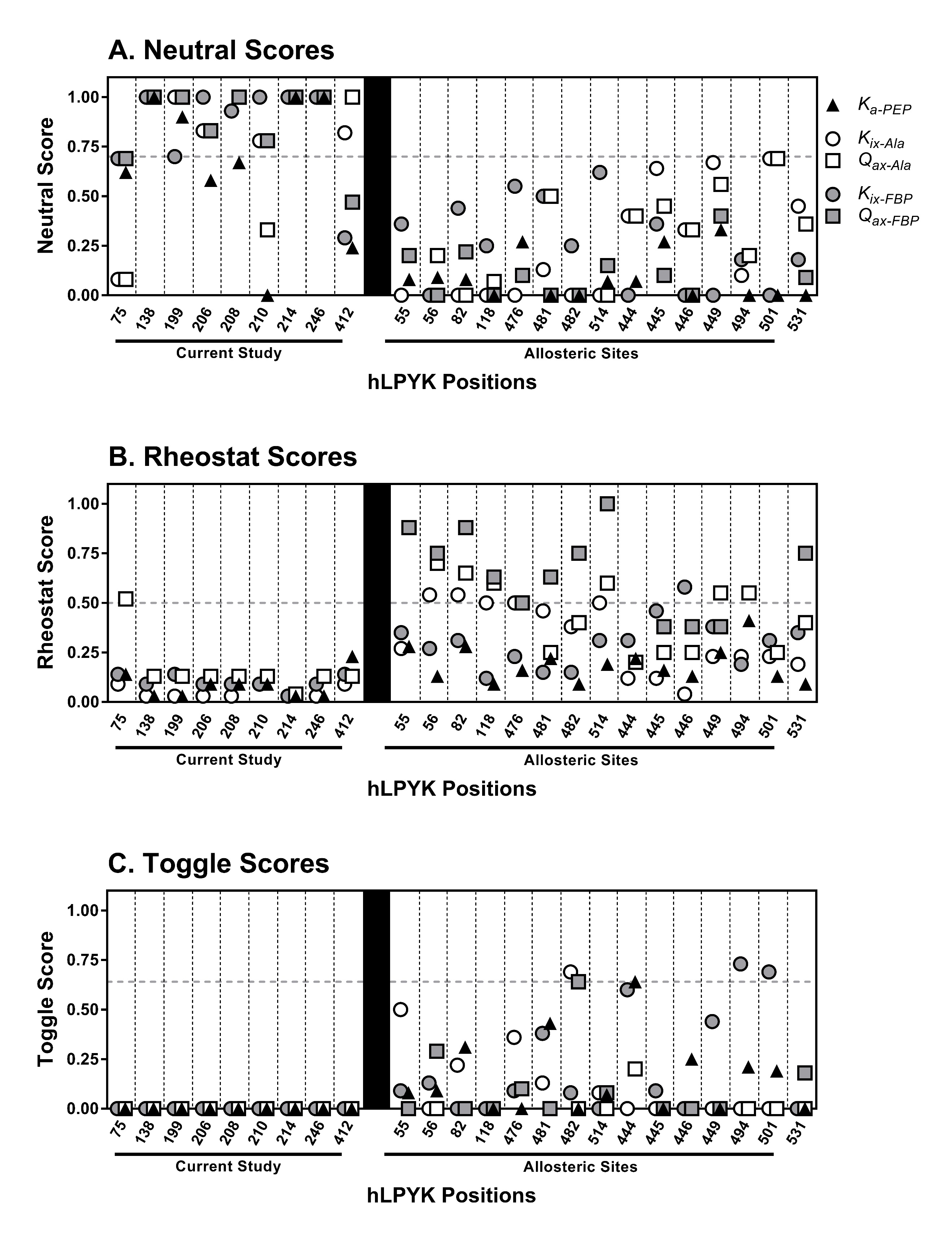

**Supplemental Figure 5**. Aggregate neutral scores for all substitutions in the two sets of positions. Results from both the current work and the previous studies of hLPYK allosteric sites ^3, 5, 27^ were evaluated as aggregate neutral scores, calculated by treating all substitutions at all positions in the respective set as a single “set” with RheoScale. The horizontal dashed line indicates the threshold of 0.7 determined from the data in Figure 5. Overall, a higher fraction of the variants in the current study had neutral outcomes than did the allosteric studies.

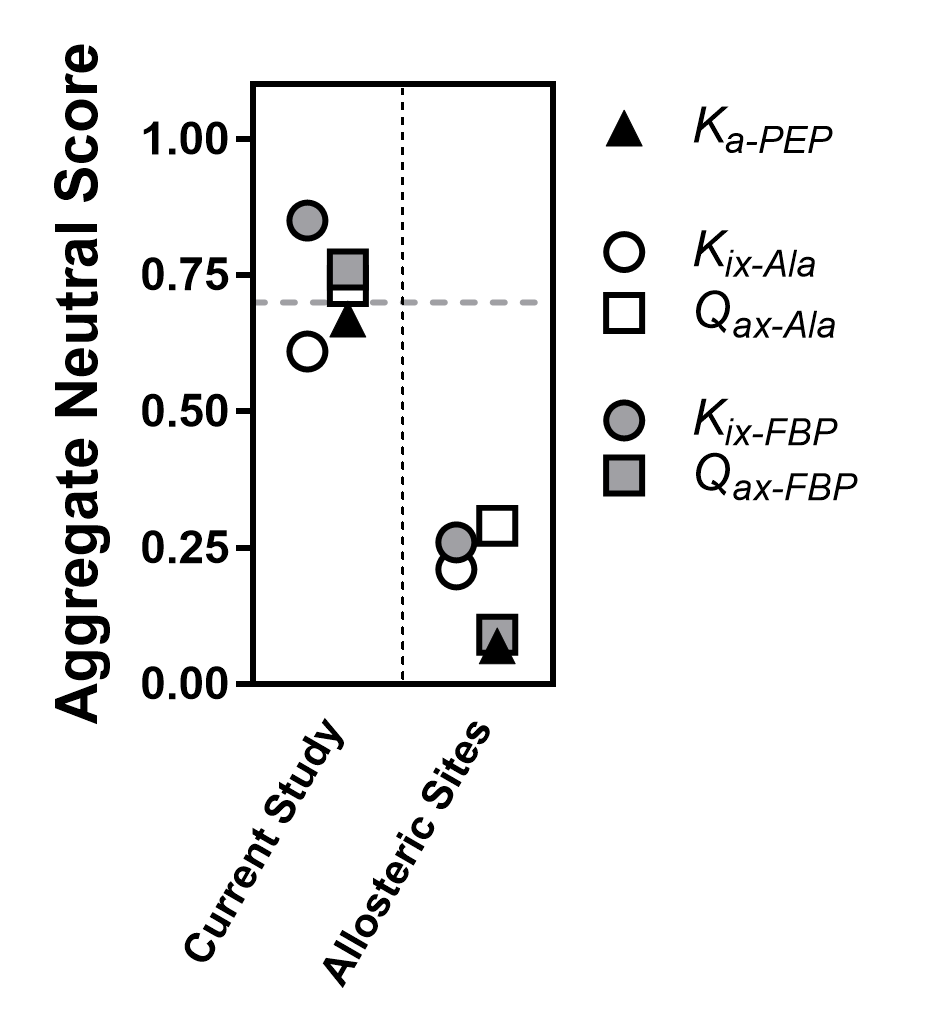

**Supplemental Figure 6. Exploration of adding weighted sub-bins to the calculations of neutral scores**. In the RheoScale analyses of our previous publication ^2^, histogram bin size was determined by dividing the range of hypothetical data by one of three criteria: (i) the minimum number of bins suggested between error, (ii) mean number of variants, or (iii) the median number of variants. The neutral score was calculated by the proportion of variants falling within the wildtype bin. This approach worked well for determining rheostat scores. However, neutral scores depend critically upon the error associated with the wildtype functional values. Therefore, in the current study, bin size was calculated by centering the bin on the average of all wildtype values and considering the Gaussian distribution expected for these data. As described in Materials and Methods, we also considered the effects of variants that fall just outside of wildtype distribution: There is some probability that these values fall on the tails of the wildtype distribution, however, some likely should fall into adjacent bins. To explore these options, five sets of adjacent bins of 1/5^th^ wildtype-bin-size were weighted 0.5, 0.4, 0.3, 0.2, and 0.1 (from the immediately-adjacent to distal bins, respectively) and included in calculations of the neutral scores. Alternative scores are graphed for each position. The addition of sub-bins had modest effects on the neutral scores. Therefore, the approach to add sub-bins was NOT used in the final analysis. However, this exercise gave us confidence that the scores in the text of Figure 5 accurately reflect the substitution behaviors of the positions examined.

**Supplemental Figure 6.**
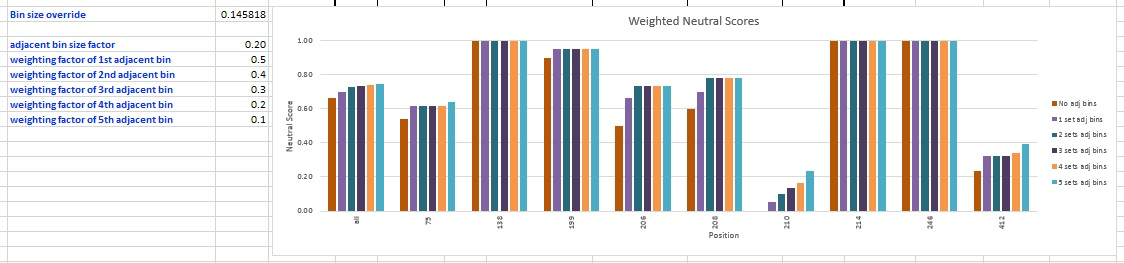
